# A Biochemical Description of Postsynaptic Plasticity – with Timescales Ranging from Milliseconds to Seconds

**DOI:** 10.1101/2023.07.09.548255

**Authors:** Guanchun Li, David W. McLaughlin, Charles S. Peskin

## Abstract

Synaptic plasticity (long term potentiation/depression (LTP/D)), is a cellular mechanism underlying learning. Two distinct types of early LTP/D (E-LTP/D), acting on very different time scales, have been observed experimentally – spike timing dependent plasticity (STDP), on time scales of tens of ms; and behavioral time scale plasticity(BTSP), on time scales of seconds. BTSP is a candidate for the mechanism for rapid learning of spatial location by hippocampal place cells. Here a computational model of the induction of E-LTP/D at a spine head of a synapse of a hippocampal pyramidal neuron is developed. The single compartment model represents two interacting biochemical pathways for the activation (phosphorylation) of the kinase (CaMKII) with a phosphatase, with Ion inflow described by NMDAR, CaV1, and Na channels. The biochemical reactions are represented by a deterministic system of differential equations. This single model captures realistic responses (temporal profiles with the differing timescales) of STDP and BTSP and their asymmetries for each (STDP or BTSP) signaling protocol. The simulations detail several mechanisms underlying both STDP and BTSP, including i) the flow of *Ca*^2+^ through NMDAR vs CaV1 channels, and ii) the origin of several time scales in the activation of CaMKII. The model also realizes a priming mechanism for E-LTP that is induced by *Ca*^2+^ flow through CaV1.3 channels. Once in the spine head, this small additional *Ca*^2+^ opens the compact state of CaMKII, placing CaMKII “in the ready” for subsequent induction of LTP.

---

Synaptic plasticity is an essential mechanism that underlies learning and memory. At present, however, synaptic plasticity, with its many diverse forms, is far from thoroughly understood. Experimental studies have shown that electrical or spiking stimuli can lead to a strengthening or weakening of synapses lasting for hours, days to years –termed long-term potentiation/depression (LTP/LTD) (1, 2). STDP (spike timing dependent plasticity) is a form of synaptic plasticity (3–5), which is induced by a standard stimulus protocol of many repetitive pairs of pre-synaptic and post-synaptic spikes, with a short time interval of tens of milliseconds separating the pre-post members of each pair. STDP is *Hebbian*, with LTP (LTD) occurring when the pre-synaptic spike precedes (follows) the post-synaptic spike of each pair (6). While STDP is a beautiful realization of Hebbian learning, there are probably other mechanisms of synaptic plasticity that underlie other types of learning. For example, rodent hippocampal CA1 place cells learn location with remarkable ease, after the rodent makes only one (or a few) passes across the specific location. The time scales and “one-shot” nature of learning are difficult to reconcile with the properties of STDP (7–9). Recently, a very different form of synaptic plasticity, BTSP (behavioral time scale plasticity), has been observed to emerge from a different stimulus protocol (10). This BTSP form of LTP seems more likely to underlie such one-shot learning, with its characteristics very different from those of STDP. BTSP operates over much slower time scales (seconds, rather than tens of milliseconds), and is not Hebbian (with LTP occurring whether or not the pre-synaptic excitation occurs before or after the post-synaptic excitation).

Experimental studies have identified two elements as vital for LTP/D. *First*, the intracellular calcium influx through NMDAR and CaV channels has been shown to be necessary and sufficient for LTP/LTD (11–14). *Second*, two proteins (CaMKII and Phosphatase) that are activated by the calcium influx are known to modify the synaptic efficacy. The activation (phosphorylation) of the kinase CaMKII is initiated by the Calcium-CaM complex and sustained by the autophosphorylation of CaMKII. This active state of CaMKII then triggers a conformational change of the AMPARs, leading to a strengthening synaptic transmission (15–17). Similarly, the phosphatase (PP), which catalyzes dephosphorylation, has also been shown to be necessary for the induction of LTD (18–20). In addition, Tsien and colleagues have argued from existing experimental results (21–25) to propose a *priming* mechanism that could enhance one-shot learning (26). This priming mechanism is induced by a small steady depolarization that causes a small concentration of Calcium to flow through CaV1.3 channels into the spine head, where it would induce a transient activation of CaMKII, placing the synapse “in the ready” for a stronger one-shot signal to induce LTP.

Our work began from some questions: Can a single computational model of the biochemical reactions within the spine head at a synapse realize both STDP and BTSP when stimulated by the STDP or BTSP protocols, respectively? If so, what are the mechanisms by which this single model realizes the very different characteristics of STDP vs. BTSP (vastly different time scales and Hebbian nature)? In addition, can a priming mechanism involving calcium channels (CaV channels) (26) be realized in the same model; and, if so, what are the characteristics of the model’s priming mechanism and how does it relate to BTSP and one-shot learning?

In this paper, we construct a realistic biological model of the post-synaptic biochemical reactions at a single synapse during early LTP/D (E-LTP/D). We start from Graupner & Brunel’s model (27), which successfully reproduces the response to a STDP protocol with a bistable single-compartment system of ordinary differential equations (ODEs). Starting from this model, we replace several components with a more fundamental and precise description; we also remove the traditional I1-PP1 signaling pathway and substitute instead a phenomenological representation from (28). The resulting model is tri-stable with three coexisting stable steady states directly related to the synapse’s basal, LTP & LTD states. Under stimulation, the system makes transitions from the basal state to either of the LTP/D states – corresponding to the occurrence of LTP/D. In reality, these states of CaMKII and PP are likely at best *meta-stable*, becoming unstable over longer time scales by mechanisms not modeled here.

The paper is organized as follows: After presenting the model, we then validate that it has a realistic response to both a prescribed calcium stimulus and a STDP signaling protocol, and that it captures the distinct behavior of LTP/LTD and the short timescales associated with the STDP protocol. We then establish that a priming picture can be realized in the model, through a mild depolarization that opens CaV1 channels to a *Ca*^2+^ inflow that primes the activation of CaMKII. This establishes the importance of CaV1-CaMKII signaling in the model; and, thus, its possible importance for one-shot learning. Next, we confirm that the model exhibits a realistic response to the BTSP protocol (the same model, with no changes in parameter values), demonstrating the model’s ability to reproduce the long timescales and the non-Hebbian nature of the BTSP. We then analyze the timescale of CaMKII activity within the model and specify the central role of CaMKII in integrating the information of the BTSP pre-synaptic and post-synaptic stimuli spaced several seconds apart, thus, showing how BTSP can share a similar seconds-long timescale with CaMKII activity. Finally, we explore how priming could i) strengthen CaMKII’s role as an information integrator and ii) extend the timescale of BTSP.

## Results

### Qualitative Description of the Model

We have developed and implemented a computational model of the biochemical reactions responsible for E-LTP/D for synapses of CA1 pyramidal neurons in the hippocampus. The reaction scheme is illustrated in Fig.1A. The presynaptic spikes, postsynaptic spikes, and plateau potential open the relevant calcium channels, allowing the calcium ions to flow into the spine head. The increasing calcium concentration will activate CaMKII and phosphatase, which will further activate themselves through autocatalytic reactions, while the two will inhibit each other through phosphorylation and dephosphorylation.

**Fig. 1.**
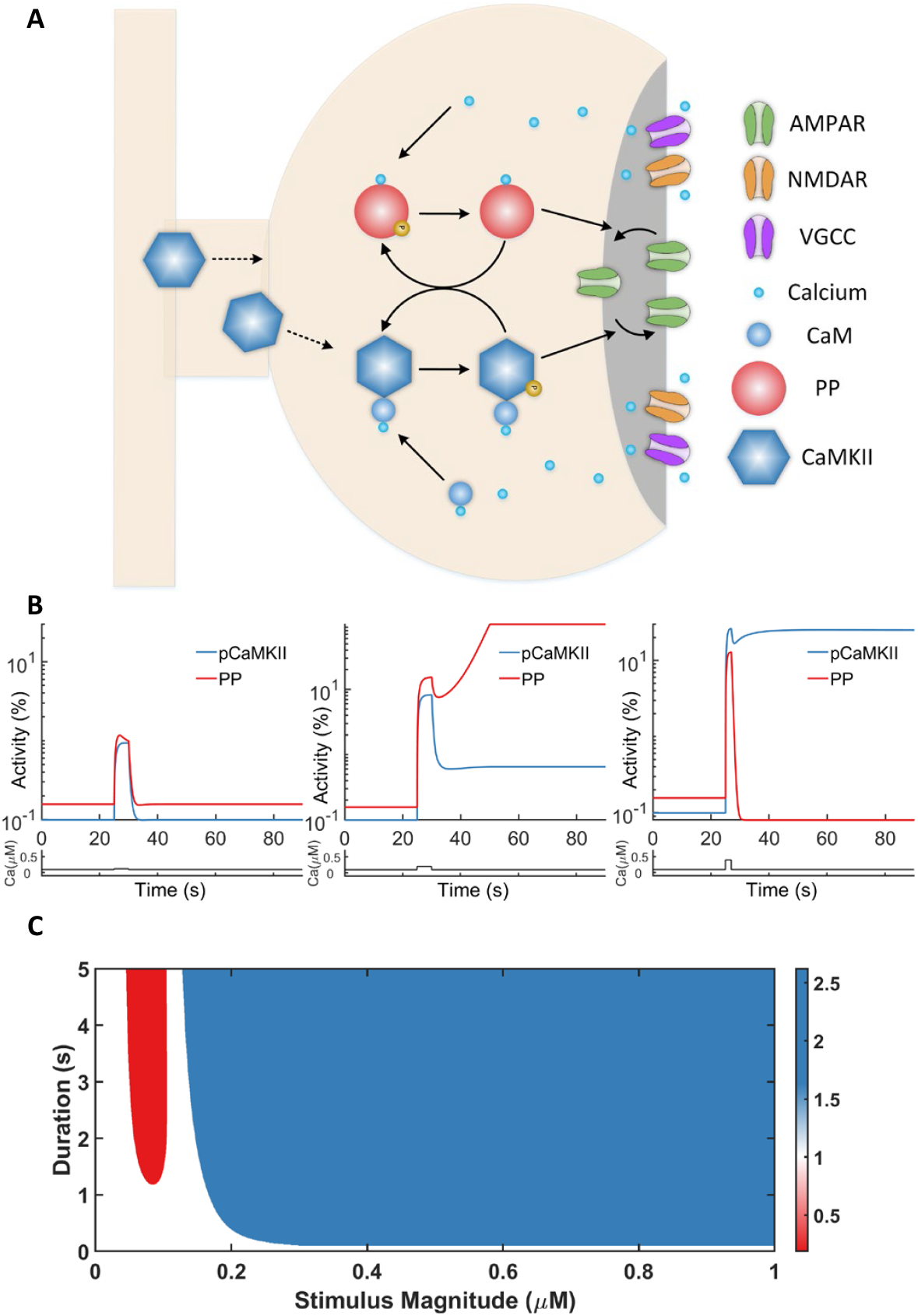
Calcium-Induced Synaptic Plasticity: A Model of CaMKII and PP Interaction. (A) Chemical reaction scheme of the model. Calcium flows into the synapse head through NMDAR and VGCC (voltage-gated calcium channels) and activates CaMKII (after binding with CaM to form the Calcium-CaM complex) and PP. Active CaMKII and PP mutually inhibit through phosphorylation and dephosphorylation. (B) pCaMKII (phosphorylated CaMKII) and active PP level during a calcium pulse stimulus. The top graphs show the phosphorylated CaMKII (blue traces) and active PP (red traces) concentration, and the calcium concentration is given at the bottom (black traces). Left panel, CaMKII, and PP are weakly activated during a weak pulse (0.02*μM*, 5 seconds) and return to the basal state when *Ca*^2+^ is removed. Middle panel, PP is activated by a slightly stronger stimulus with the same duration (0.1*μM*, 5 seconds) and stays active, leading to LTD. Right panel, During high level *Ca*^2+^ stimulus (0.3*μM*, 2 seconds), both CaMKII and PP are active while the kinase activity is dominant after the stimulus. (C) Dependence of the sign of synaptic modification on *Ca*^2+^ level and duration during a pulse (Note that for any fixed duration, the results are similar to the BCM rule).

The model describes how CaMKII and phosphatase activities respond to intracellular calcium dynamics determined by presynaptic and postsynaptic electrical events. It is a single compartment model of the post-synaptic density, containing well-mixed *Ca*^2+^ ions, CaM, CaMKII, and phosphatase. The biochemical reactions of the two interacting pathways [the kinase CaMKII pathway and a phenomenological phosphatase PP pathway] are described by a deterministic system of ordinary differential equations. Ion channels, such as NMDAR, CaV1, and Na channels, describe the ion inflow (including *Ca*^2+^). The dynamics exhibit three stable fixed points, representing basal, LTP, and LTD states. Importantly, the model’s transient dynamics describes the phosphorylation of CaMKII as the formation of E-LTP, and the dephosphorylation of phosphatase as the formation of E-LTD. The full model is summarized in the “Methods” section, with a complete detailed description in the supporting information.

The model captures the initiation of Early LTP/D, with this initial transient dynamics organized by the model’s stable *fixed points* or steady states. (Most likely, over a longer time scale, these states would be, at best metastable, being destabilized by mechanisms that are not modeled here; however, these metastable states would still organize the initial transient dynamics.) We first show that the model operates in a tri-stable regime with three coexisting stable states (the basal state, an LTP state, and an LTD state). Synaptic efficacy increases when CaMKII sustains a high activity and decreases when PP sustains a high activity.

### A tri-stable model of synapse efficiency

It is well known that intracellular calcium dynamics can induce both LTP and LTD states (12–14) Moderate, long-lasting calcium will lead to the LTD state, while a large calcium inflow will trigger LTP (29, 30). We first study our CaMKII and phosphatase system through a prescribed calcium stimulus pulse in Fig.1B. Initialized in the basal state (of low CaMKII and phosphatase activities), a large calcium pulse (0.3*μM*, for 2 seconds) will place the system within the basin of attraction of the LTP state (of high CaMKII phosphorylation and low phosphatase activity). In contrast, again beginning from the basal state, a longer calcium stimulus of moderate strength (0.1*μM*, for 5 seconds) places the system within the basin of attraction of the LTD state (of low CaMKII phosphorylation and high phosphatase activity). As a comparison, after a weak calcium stimulus (0.02*μM*, for 5 seconds) on the basal state, the system will return to the basal state, showing the basal state is stable under perturbations. The LTP and LTD states are also stable under similar perturbation experiments (Fig.S9).

To further study how the magnitudes and durations of the calcium dynamics determine the steady states, we simulate the system with different calcium pulse profiles. The results are summarized in the phase diagram of Fig.1C. In this figure, the system always starts from the basal state, similar to the above simulations. A mild stimulus, either of small magnitude or short duration, will be unable to push the system outside of the basin of attraction of the basal state. However, a stimulus with a moderate magnitude between 0.075 *−* 0.125*μM* will induce the system to an LTD state when it is of sufficient duration (larger than 1 second). A strong stimulus with a magnitude larger than 0.2*μM* will lead to LTP. It is noteworthy in the phase diagram that a more extended time window is required for LTD to happen, which is consistent with experimental observations (29, 30). We also observe an intermediate region in Fig.1C for the basal state between LTD and LTP (when the stimulus magnitude is around 0.15*μM*). This intermediate state is similar to that observed in the phenomenological model of (28), Fig.2D, where the authors argue a relationship with the switch from depression to potentiation in the BCM rule (31).

**Fig. 2.**
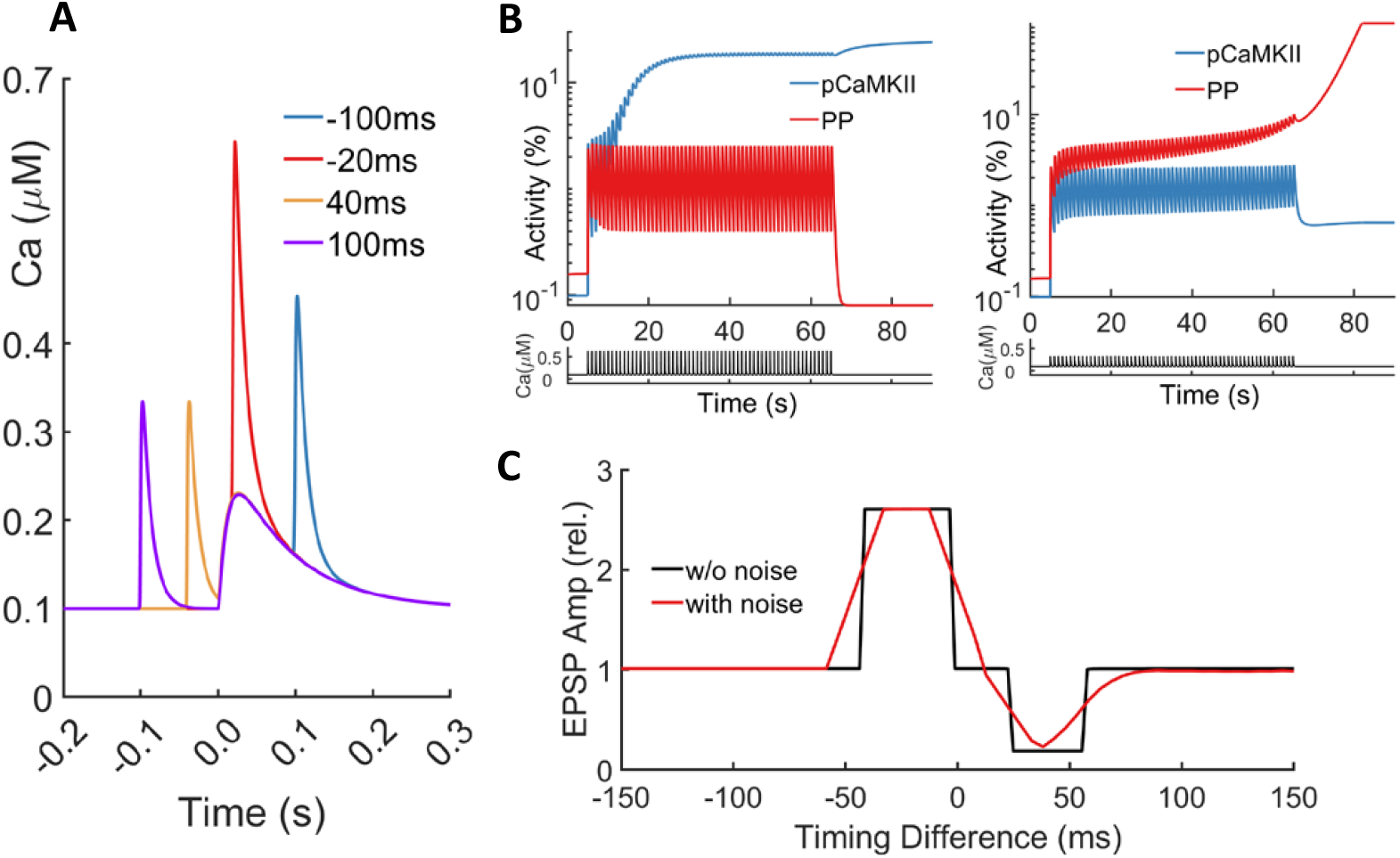
Synaptic Modification in Response to STDP protocols. (A) Calcium dynamics during a pair of a pre-synaptic and a post-synaptic spike at different timing differences Δ*t*. The time course of the intracellular calcium concentration is generated by the model in response to a pre-synaptic spike at *t*_pre_ = 0 ms and a post-synaptic spike at *t*_post_ = Δ*t* ms. The Δ*t* values are noted in the legend. (B) Temporal evolution of Calcium, PP1 activity, and phosphorylated CaMKII activity level during STDP stimulation protocols with Δ*t* = *−*20 ms (left panel) and Δ*t* = 40 ms (right panel). 60 pairs of pre-synaptic and post-synaptic spikes in 1Hz evoke LTP at Δ*t* = *−*20 ms but lead to LTD at Δ*t* = 40 ms. (C) Synaptic modifications in response to STDP protocols. In the deterministic (black curve) setting, STDP protocols with Δ*t* between *−*40 and 0 ms evoke LTP, and Δ*t* between 20 and 60 ms evoke LTD. In the stochastic setting (red curve), STDP protocols with Δ*t* between *−*55 and 10 ms evoke LTP, and Δ*t* between 10 and 85 ms evoke LTD.

It is known experimentally that both LTP and LTD can be reversed with appropriate stimuli, which is termed depotentiation and dedepression, respectively (32, 33). We conduct simulations similar to those in (28) and present the results in Fig.S5B. A moderate pulse stimulus (0.08*μM*, 10 seconds) reverses LTP, moving it back to the basal state, while a substantial calcium influx (0.35*μM*, 3 seconds) can reverse the LTD to the basal state.

### STDP simulations

We have just described the responses of the model to prescribed calcium inflow. However, in experiments calcium inflow is not directly prescribed; rather, the experiment controls synaptic events (pre-synaptic and post-synaptic spike trains and dendritic plateau potentials) through current injection, which in turn leads to calcium inflow through NMDA and CaV channels. Following (27), we simulate the calcium inflow that results from these prescribed synaptic events, as described in detail in Methods and Supplementary Material.

LTP induced by spike timing-dependent plasticity (STDP) has been observed in many experiments (see for example (3–5)), with the traditional signaling protocol that of (5). Pairs of pre-synaptic and post-synaptic spikes are evoked repetitively with controlled timing differences between the two spikes. The synapse is strengthened when the pre-synaptic spikes precede the post-synaptic ones, while it is weakened if the post-synaptic neuron spikes earlier, all within 50-100 ms. The causality relation property makes STDP a natural candidate for Hebbian learning (34, 35).

We use a standard STDP protocol containing 60 repetitive pairs of pre-synaptic and post-synaptic spikes, with a frequency of 1Hz (see (5)). The calcium dynamics evoked by one pair of spikes are shown in Fig.2A. When the pre-synaptic spike slightly precedes the post-synaptic spike, a large calcium influx results, as the magnesium block of the NMDAR channels is removed by the membrane depolarization through back-propagating action potential (bAP) of the post-synaptic neuron. When the pre-synaptic spike is slightly after the post-synaptic spike, the removal of the magnesium block is not complete, resulting in a relatively moderate increase in calcium concentration. When the two spikes occur with a time difference longer than 100ms, either positive or negative, the weak calcium inflow is almost a linear summation.

In Fig.2B we show how the response of the CaMKII and PP interacting systems to this STDP signaling protocol. For small negative dt (−20ms), the large calcium influx will activate CaMKII significantly, and the activation is accumulated steadily during each pair of spikes. In contrast, PP is activated during spikes but quickly drops back to the resting concentration when the stimulus is off, and no cumulative effect can be observed. That leads to a significant amount of activated CaMKII with no change for PP, leading to the steady state in which CaMKII dominates. For extensive negative dt (−100ms), both CaMKII and PP show the *increase-and-decay* behavior, and neither can be accumulated during the presentation of multiple pairs (Fig.S6).

Similar behavior can be discovered for positive dt, as also presented in Fig.2B. For small positive dt (40ms), the long, moderate calcium influx will activate PP, and the activation is accumulated during each pair of stimuli. At the same time, it will only induce a moderate activation of CaMKII, further restricted by the significant inhibition by active PP, which places the system in the attractor basin of the LTD state. For large positive dt (100ms), although both CaMKII and PP accumulate slightly, the accumulation does not lead the system to either LTP or LTD state (Fig.S6). Thus, the different calcium inflow profiles lead to the particular activity of CaMKII and PP, which finally leads to the asymmetric STDP.

The system’s response under the STDP protocol for a broader range of the timing difference dt (the pre-synaptic spike time minus the post-synaptic spike time, within one pair) is shown in Fig.2C. We notice that LTP occurs for a short interval of negative dt, while LTD is induced for a somewhat longer interval of positive dt, both in tens of milliseconds. The system will return to the original steady state for timing differences beyond that range. Besides, between the LTP interval and LTD interval, there is an intermediate area near Δ*t* = 0 ms where no learning happens, which is also seen in other simulation studies (27, 36).

The stochastic opening of ion channels, a primary contributor to the noise of calcium dynamics, will smooth these deterministic results. To represent this stochastic effect, we include two sources of noise following (27): (i) random maximum conductance of NMDAR at each presynaptic spike and (ii) random maximum conductance of the CaV channels at each postsynaptic spike. Similar to the deterministic case, the average of AMPARs’ number is used as the indicator of synaptic efficacy, also related to the transition probability from the basal state to the LTP or LTD state. The smoothed results are also shown in Fig.2C. The synaptic efficacy is approximately an exponentially decaying function of the timing difference *dt* for both positive and negative intervals. The result is consistent with experimental observations and previous numerical simulations, in particular, the asymmetric behavior of LTP/LTD and the short-range learning window over several tens of milliseconds.

### Voltage-dependent learning

Experiments have shown that place cells can be firmly induced in vivo by the injection of a small direct-current (DC) stimulus, which only increases the baseline membrane potential by several millivolts (24). This induction is mainly reversible, and has been shown to result from the voltage dependency of peri-somatic channels (37). Here we investigate the model’s LTP dependence on the baseline membrane potential, which could be related to the small residual irreversible components observed by (24). It is not clear how the place cell is induced in vivo; thus, there are possibly several different experimental protocols that can lead to induction. Here we borrow from an in vitro protocol used in (10) in a study of the production of LTP and the formation of place cells. Our stimulus protocol consists in a plateau potential (which can be induced by sodium current injection or naturally occur (10)), following three pre-synaptic spikes (versus ten in (10)), together with a small DC mimicking the mild depolarization used in (24).

The calcium dynamics under our protocol (with and without DC injection) is shown in Fig.3A. Given that the timing difference between the pre-synaptic spikes and plateau potential is large enough (500ms), the calcium dynamics is a superimposition of the two stimuli with no overlapping effects. The DC injection increases the resting membrane potential, which increases the baseline calcium concentration by 5% (from 0.1 muM to 0.105 muM) through the opening of CaV1.3 channels. In addition, it also slightly releases the magnesium block of the NMDA channels, allowing a significant calcium influx during pre-synaptic spike train.

**Fig. 3.**
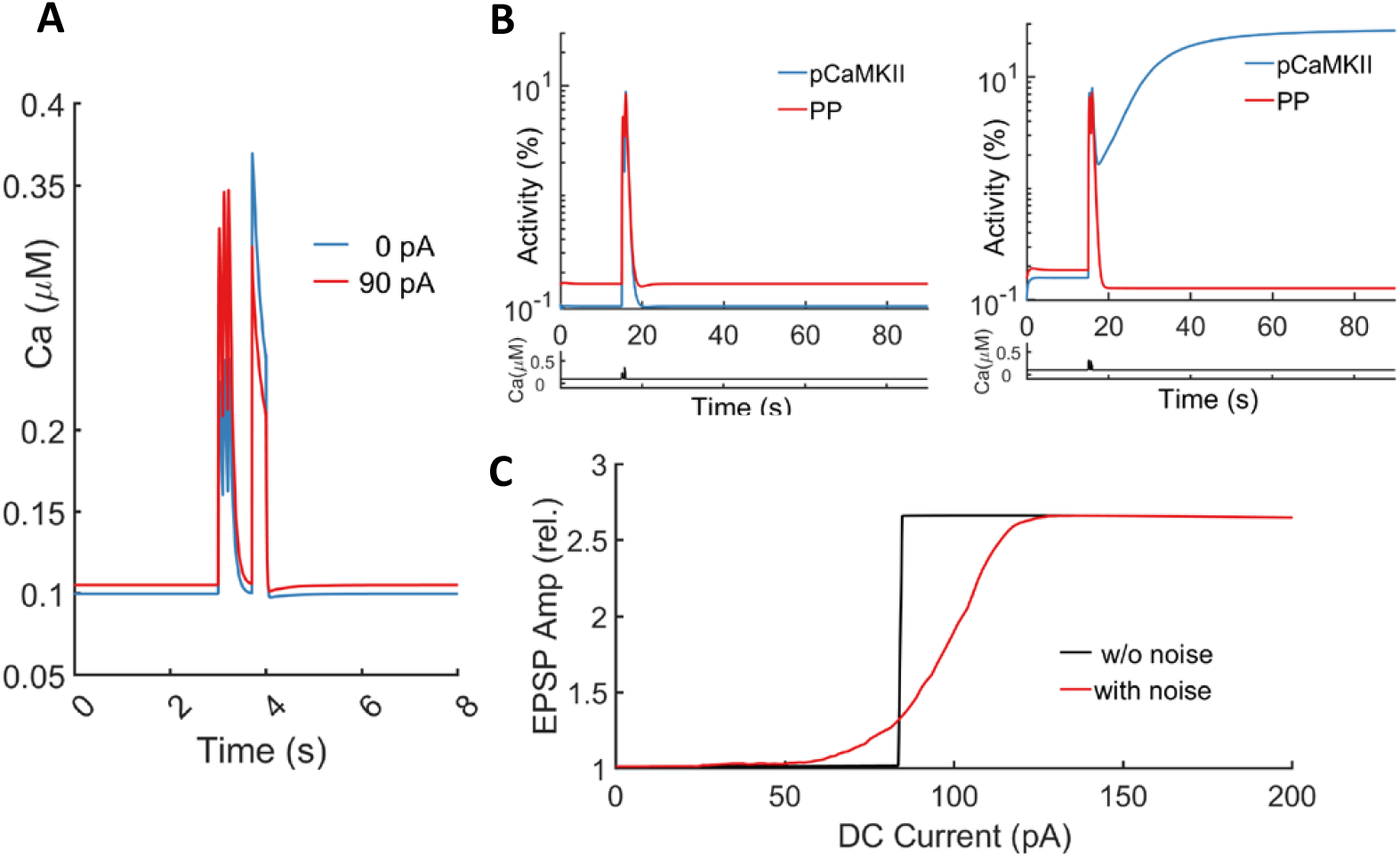
Synaptic Modification in Response to Voltage-Dependent Learning. (A) Calcium dynamics during the combination of a plateau potential and pre-synaptic spike trains, with and without DC stimulus. The time course of the intracellular calcium concentration is generated by the model in response to three pre-synaptic spikes starting at *t*_pre_ = 3 second in 10Hz and a 300 ms plateau potential initiating at *t*_post_ = 3.7 ms. The 90pA DC increases the baseline calcium concentration from 0.1*μM* to 0.105*μM*. (B) Temporal evolution of Calcium, PP1, and phosphorylated CaMKII activity level during the voltage-dependent one-shot learning. The left panel shows the results with DC, and the right panel shows the results without. No LTP is induced without DC, while a sustained 100pA current injection is enough to switch the response to CaMKII domination (LTP). (C) Synaptic modifications in response to voltage-dependent one-shot learning with varying amounts of DC injection. A sharp transition to learning occurs at 85pA in the deterministic setting (black curve), and a similar transition is observed in the range of 70-130pA for the stochastic setting (with the half height of the transition at 100pA).

The results with and without DC are shown in Fig.3B. For the case without DC, we see mild increases of CaMKII and PP during the two stimuli, with PP slightly dominating the competition. The system rapidly decays back to the basal state. For the case with DC, a larger amount of CaMKII is activated during the pre-synaptic spikes, and is further amplified by the persistent, significant calcium influx induced by the plateau potential. The phosphorylated CaMKII dominates after the stimulus and dramatically inhibits the activity of PP, which finally leads to LTP. The above results show how several millivolts’ increase of membrane potential could switch the system from no learning to learning. It is worth emphasizing that during this shallow voltage range, the voltage dependency of NMDAR channels is mild; in contrast, CaV channels (especially CaV1.3) have a much steeper voltage dependency [Fig.S8A]. As a result, the CaV channels contribute to most of the changes in calcium dynamics by DC injection when there are no pre-synaptic spikes [Fig.S8B].

Next, we investigate how the amplitude of the DC injection will influence learning, by injecting DCs of different magnitudes [Fig.3C]. In all cases, the system terminates in either the basal or the LTP state, with no LTD state observed. In this deterministic setting, the transition from basal to LTP is sharp at DC = 85pA, corresponding to a baseline membrane potential of -62.5mV. When the calcium inflow is stochastic, this sharp transition is smoothed [Fig.3C]; however, it remains a steeply monotonic increasing function of the amount of DC, with a sharp transition from no learning to complete learning in the range of 70 - 130 pA, which corresponds to a shallow range (5mV) of baseline membrane voltage from -65 to -60 mV, also reported in (24) (see also Fig.S7).

### Essence of priming

Recently, a *priming* mechanism has been proposed by the Tsien Lab (26) for a possible role to enhance one-shot learning of CA3 to CA1 synapses of place cells in the rat hippocampus, based on a detailed inspection of experimental results about CaMKII activity. This priming picture can be described as follows: The synapse can get primed by mild depolarization and eventually become potentiated if a plateau potential follows within a few seconds. This primed state itself is not stable, as the synapse can return to its original, non-primed state after the mild depolarization is turned off. An intermediate, partially active state of CaMKII serves as a priming for the following stimulus. The mild depolarization required for priming lies in a voltage range where voltage-dependent CaV channels (CaV1.3) show steep dependency, indicating a possible important role for CaV channels in hippocampal place field formation.

As shown in the previous section, DC injection can influence learning. Here we make a direct detailed comparison of the dynamics with and without DC. [Fig.4A]. Before the onset of pre-synaptic spikes, the most observable difference is the increase of *oCaMKII* (open but not phosphorylated) CaMKII caused by the application of DC. The more open CaMKII, combined with the greater calcium influx during pre-synaptic spikes, prepares the system (more oCaMKII and *pCaMKII* (phosphorylated CaMKII)) before the onset of plateau potential – making it easier for the plateau potential to induce LTP, which is exactly the meaning of priming. In addition, this priming preparation is a transient activity of CaMKII, which can be reversed when no following stimulus is presented, and DC is removed – also consistent with the meaning of priming. Conclusively in the computational model, the transient activity is a priming as it satisfies i) transient activation from mild depolarization that ii) helps the learning.

**Fig. 4.**
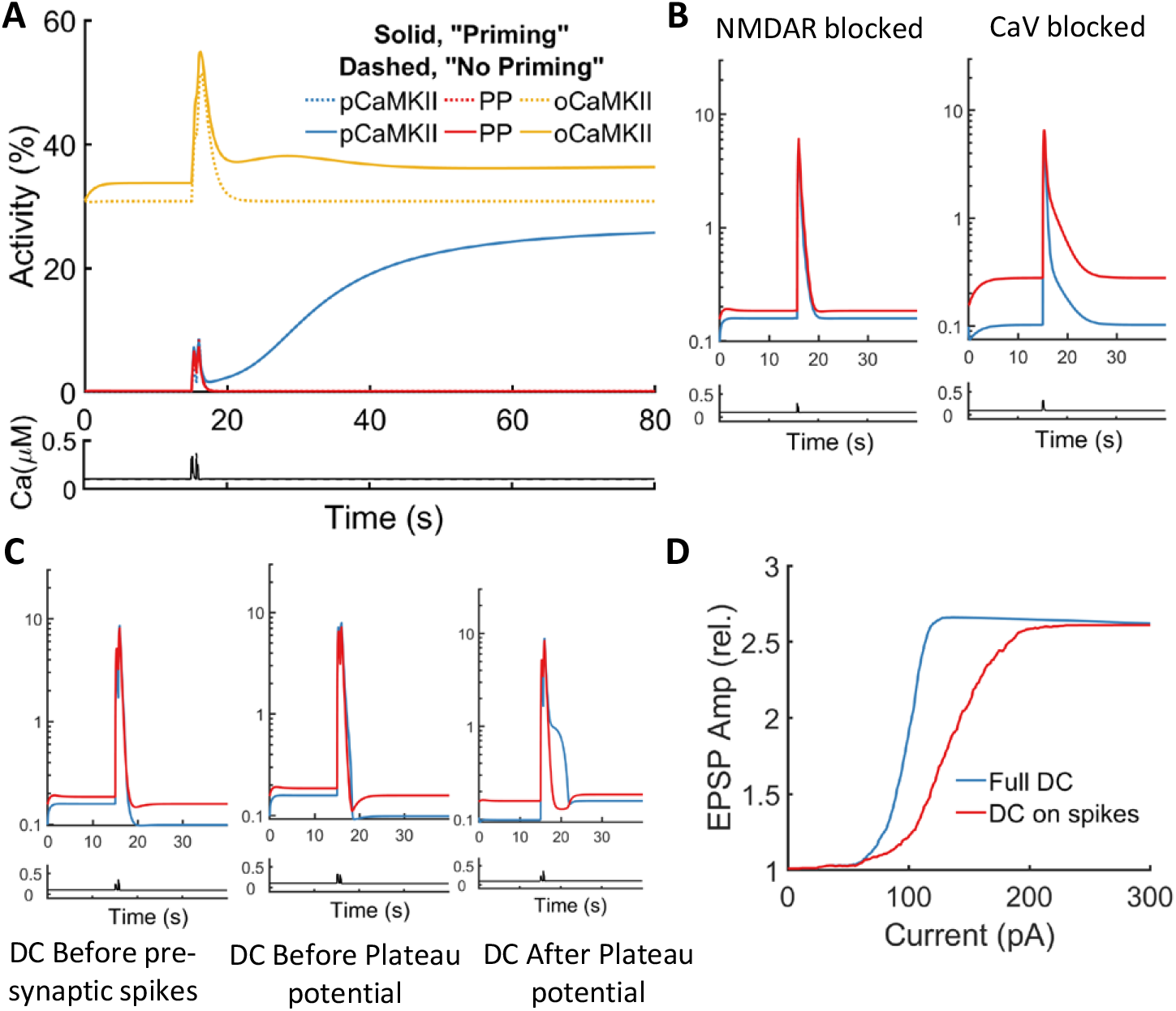
Priming and the Role of NMDAR and CaV Channels. (A) Comparison of time courses of calcium, PP1, phosphorylated, and open CaMKII activity level during the one-shot learning protocol, with and without DC (priming). The intracellular calcium increases by 5% with DC, raising open CaMKII by 10% (same data as in Fig.3B, different scales). (B) Temporal evolution of Calcium, PP1, and phosphorylated CaMKII activity level during the one-shot learning protocol, with NMDAR or CaV channels blocked. Blocking NMDAR channels (left panel) or CaV channels (right panel) abolishes LTP. (C) Temporal evolution of Calcium, PP1, and phosphorylated CaMKII activity level during the one-shot learning protocol, with transient DC. Left panel, activities with DC only before pre-synaptic spike trains. Middle panel, activities with DC only before plateau potential. Right panel, activities with DC only after plateau potential. None of the three stimuli is enough to induce LTP. (D) Synaptic modifications in response to voltage-dependent one-shot learning with varying amounts of DC injection in the stochastic setting. The black curve shows the results with DC throughout the experiment (same data as Fig.3C), and the red curve shows the results with DC applied only during pre-synaptic spike trains. Transitions are observed at both curves, but a higher range of DC (90 - 210 pA, half height at 150pA) is required for the transition to happen with depolarization only on pre-synaptic spikes compared with complete DC (transition occurs at 70-130 pA, half height at 100 pA).

Next, we describe experiments with different channels shut down to investigate the role of ion channel types in the above protocol and priming. Here, we consider the results for NMDAR and CaV channels. As shown in Fig.4B, shutting down NMDAR will prevent calcium inflow during pre-synaptic spikes, weakening the priming and preventing learning. Blocking CaV channels, alternatively, will abort the calcium flow during the plateau potential, leading to no learning. These effects within the model of blocking ion channels are consistent with experimental results (10), in which either blocking NMDAR channels or CaV channels will prevent the emergence of place cells.

Although we have shown above that both NMDAR and CaV are essential for learning under the above protocol, it remains to describe the roles they play in priming and its relationship to the mild depolarization from DC injection. First, the CaV channels must play an essential role in priming, as they are the only channels influenced by DC before pre-synaptic spikes and after the plateau potential [Fig.S8B]. Next, we look at the three stages highly influenced by DC: before pre-synaptic spikes, during spikes, and after plateau potential. (We neglect the stage “during plateau potential” since the DC contribution is negligible compared with the current used to generate the plateau potential during that period.) As shown in Fig.4C, DC in any of the three stages is not enough to support LTP. Their mechanisms work together to help the learning.

The DC injection can increase the baseline calcium concentration in two ways: i)through CaV channels and ii) through NMDAR channels during pre-synaptic spikes (as it increases the baseline membrane potential). We compare the two cases (DC throughout the experiment and DC only during pre-synaptic spikes) to see how increased baseline calcium concentration helps the learning in more detail [Fig.4D]. Although it is still possible to induce LTP with DC injection only during pre-synaptic spikes, a larger amount of DC is required than in the “full-time” DC. We suggest that if there is no increase to the baseline calcium concentration during the priming stage, a stronger stimulus must be presented to induce LTP, for example, by either a prolonged plateau potential or by a larger calcium influx through NMDAR Channels with magnesium block removed more thoroughly. The last could be done with a DC injection much larger than the one to activate CaV channels.

### BTSP protocol reproduced

BTSP (Behavioral timescale plasticity) is a recently observed plateau-potential induced plasticity rule in CA1 pyramidal neurons (10, 38), with characteristics that make it a likely candidate for the synaptic plasticity underlying types of one-shot learning. Unlike classical Hebbian learning, the synapse will be potentiated if the corresponding pre-synaptic neuron fires seconds before or after the plateau potential, and over a much longer time separation than STDP. BTSP does not require 20 presentations of stimulus pairs to induce LTP; rather, in BTSP five or fewer repetitive stimulus pairs will induce LTP. These two features (longer time duration and much fewer repetitions) make BTSP a good candidate for the underlying mechanism for the emergence of place cells in hippocampal area CA1.

Here we describe the model’s response to a BTSP stimulus protocol. We use a stimulus protocol that is similar to one of in-vitro experiments (10) – a combination of 10 pre-synaptic spikes at 10 Hz, together with a 300ms plateau potential generated by sodium current injection, separated with a given timing difference spanning from -6000 to 6000 milliseconds. Slightly different from the experiment, a small DC current (42pA) is applied constantly throughout the simulation, increasing the resting membrane potential to -66mV. The calcium dynamics under this protocol are shown in Fig.5A. Here we use only two pairs of stimuli with a 15 second interval, as it is reported in (10) that one or two pairs can already induce LTP. As shown in Fig.5A, when the absolute timing difference dt is smaller than 1000ms, there will be an overlap between the pre-synaptic spike train and the plateau potential, which leads to a large calcium influx through the NMDAR channel. However, beyond that time duration, the nonlinear effect is not observed, and the calcium dynamics is only a superposition of the calcium responses to the two separated synaptic events.

**Fig. 5.**
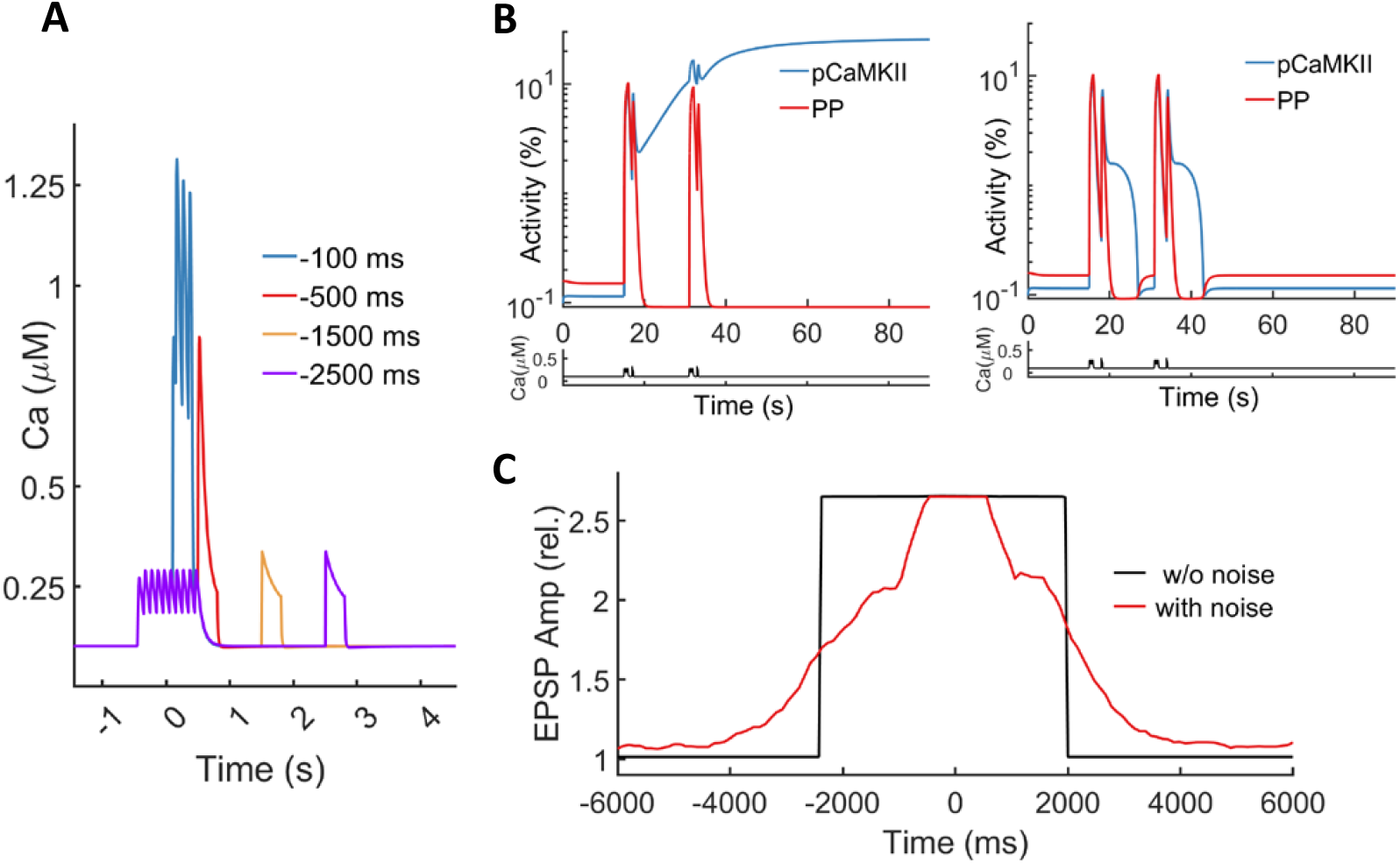
Synaptic Modification in Response to BTSP protocols. (A) Calcium dynamics during a pair of pre-synaptic spike trains and a plateau potential at different timing differences Δ*t*. It shows the time course of intracellular calcium concentration in response to ten pre-synaptic spikes at a frequency of 10Hz, centered at 0 seconds, and a 300ms plateau potential beginning at *t*_post_ = *−*Δ*t*ms, with a sustained depolarization by 42pA DC injection. The specific values of Δ*t* are noted in the legend. (B) Temporal evolution of Calcium, PP1 activity, and phosphorylated CaMKII activity level during BTSP stimulation protocols with Δ*t* = *−*1500 ms (left panel) and Δ*t* = *−*2500 ms (right panel). 2 pairs of pre-synaptic spike trains and a plateau potential (15s interval) evoke LTP at Δ*t* = *−*1500ms but have no effect at Δ*t* = *−*2500ms. (C) Synaptic modifications in response to BTSP protocols. In the deterministic (black curve) setting, BTSP protocols with Δ*t* between *−*2400 and 1950 ms evoke LTP, and Δ*t* beyond that leads to no learning. In the stochastic setting (red curve), BTSP protocols with Δ*t* between *−*4500 and 4000 ms evoke LTP, and Δ*t* beyond that leads to almost no change.

Fig.5B (and Fig.S1A) present the model dynamics of CaMKII and PP under different *dt*, the timing difference between the pre-synaptic spike trains and the post-synaptic plateau potential. For a small dt (−100ms) within the overlapping interval, the large calcium influx leads to a dramatic increase of active or phosphorylated CaMKII (pCaMKII,) which eventually causes LTP. The responses change as dt is increased in magnitude (Fig.S1A). For dt=-1500ms, one pair of the stimulus leads to a relatively minor increase of pCaMKII, but still greater than PP. A significant proportion of phosphorylated CaMKII is retained until the onset of the next pair of stimuli, when it gets amplified and dominates, producing LTP. However, if the dt is even larger (−2500ms), a single stimulus will only lead to a mild increase of CaMKII activity, which will completely diminish during the period between the two pairs of stimuli, resulting in no change in synapse efficacy. Results for positive dt are similar (Fig.S1A) and share the same dependency on the timing differences. However, in this case of positive dt, the range of dt which can induce LTP is smaller than it is for negative dt, an asymmetry that we will discuss below.

Fig.5C shows BTSP for a wide range of dt. In particular, we note that even if the two signals are separated by several seconds, the synapse is still be steadily potentiated – a long timescale consistent with experiments. The model’s BTSP results when the calcium inflow is stochastic are also presented. In this case, the timescales remain several seconds, as does the asymmetry between “pre-post” and “post-pre”, with the time range for potentiation longer for “pre-post” than for “post-pre”, exactly as reported in (10). Finally, we emphasize the role in the model of the small DC that increases the baseline membrane potential: Without it, the CaMKII in the model still gets potentiated; however with LTP only occurring for shorter dt than with the DC [Fig.S1B]. The small DC makes it easier for the model to perform consistently with experimental observations. We note that (39) observe that increasing membrane excitability helps the expansion of the timescale of BTSP, consistent with the model’s performance.

### Mechanism underlying long timescale

It is noteworthy that BTSP facilitates a seconds-long timing difference between pre-synaptic and post-synaptic electrical events, several tens times larger than that of STDP, which is typically smaller than 100 milliseconds. It is well known that the calcium dynamics inside the synapse decay very rapidly due to the fast diffusion of calcium ions. Besides, our simulation results indicate that the NMDAR channels, which play the role of coincidence detectors in STDP, cannot detect the pre-synaptic signal and post-synaptic signal together if they are placed in a timing difference larger than 300 milliseconds, consistent with the timescale of NMDAR activity reported experimentally (40). Both of these properties suggest that neither the NMDAR channels nor the calcium dynamics are suitable candidates to capture the timing difference between pre-synaptic spikes and plateau potential.

To find out what contributes to the long timescale of BTSP in the model, we first trace the activities of CaMKII and phosphatase and the relationship between the stimulus and these activities. The typical decay of open CaMKII, phosphorylated CaMKII, and active phosphatase after pre-synaptic spike trains is presented in Fig.6A. As the system is not perturbed outside of the attractor basin of the basal state, all three will decay back to the resting concentration if no extra stimulus follows, in a manner close to exponential decay, as shown in the figure. It is apparent in Fig.6A that the decay of oCaMKII is much slower compared to the decay of pCaMKII and phosphatase. The decay of oCaMKII mainly depends on the rates of docking and undocking of CaMKII subunits, while the decay of pCaMKII and phosphatase is due to their mutual inhibition, which can be quite rapid when both are highly active.

**Fig. 6.**
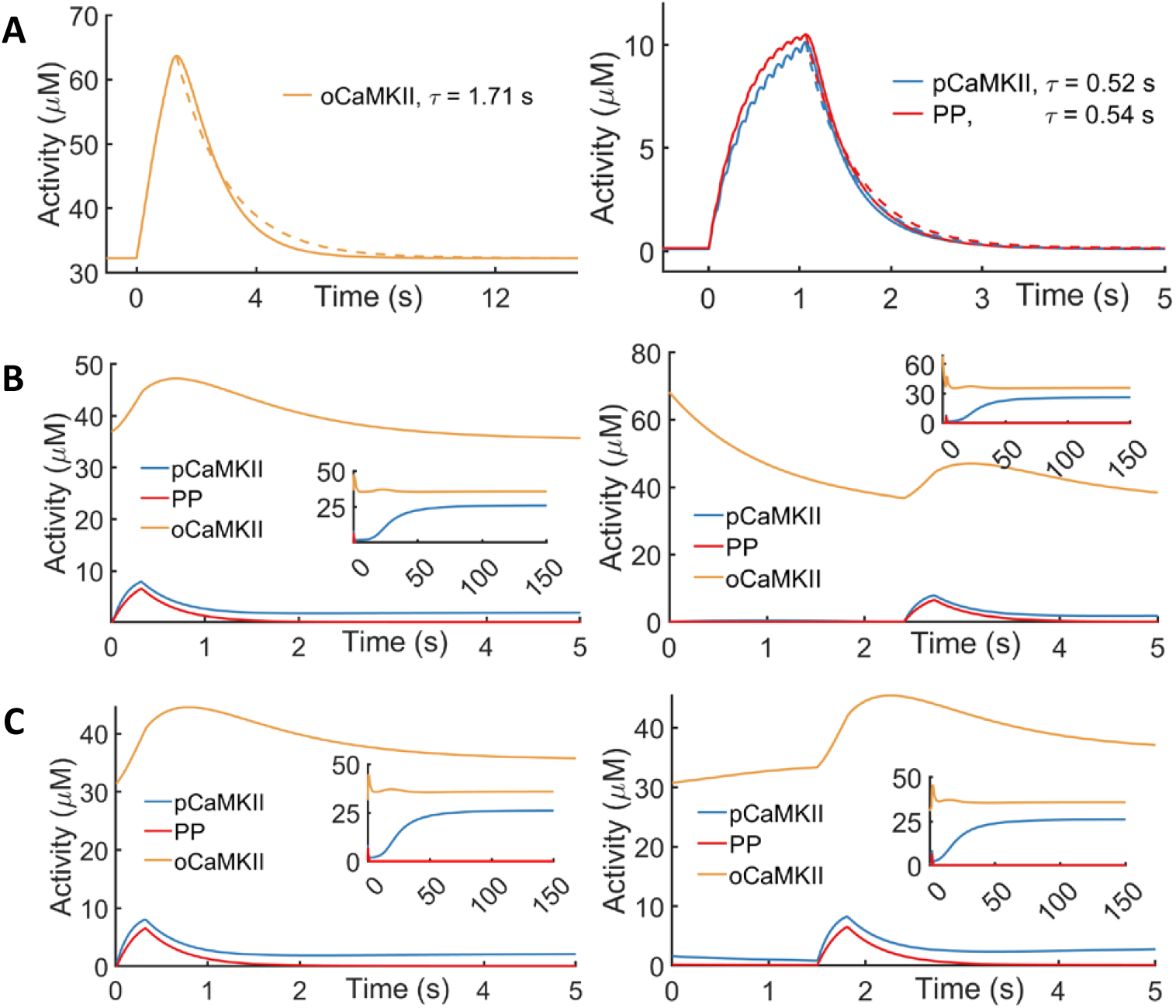
Mechanism underlying long timescale. (A) Estimating timescales of open CaMKII, phosphorylated CaMKII, and PP activity. The decay dynamics of open CaMKII (left panel), phosphorylated CaMKII, and PP (right panel) after ten 10Hz pre-synaptic spikes starting at 0 seconds are shown. All the decay dynamics can be fitted well with an exponential decay (dashed line), with time constants noted in the legends. (B) Temporal evolution of PP1, phosphorylated and open CaMKII activity level in response to a plateau potential with a perturbation on open CaMKII activity. The plateau potential is initiated by a 5000pA current injection, lasting for 300ms, starting at *t* = 0 second (left panel) or *t* = 2.4 second (right panel), accompanied by a sustained 42pA depolarization. A perturbation that increases oCaMKII by 4.8*μM* (12%) at the beginning of plateau potential (left panel) or increases oCaMKII by 32*μM* (100%) 2.4 seconds before plateau potential (right panel) evokes LTP (as shown in the insets). Neither scenario evokes LTP when the perturbation is not presented. (C) Temporal evolution of PP1, phosphorylated and open CaMKII activity level in response to a plateau potential with a perturbation on phosphorylated CaMKII activity. The plateau potential is initiated by a 5000pA current injection, lasting for 300ms, starting at *t* = 0 second (left panel) or *t* = 1.5 second (right panel). A perturbation that increases pCaMKII by 0.75*μM* at the beginning of plateau potential (left panel) or increases pCaMKII by 1.5*μM* 1.5 seconds before plateau potential (right panel) evokes LTP (as shown in the insets). Neither scenario evokes LTP when the perturbation is not presented.

Although we have shown in the previous sections that the activity of oCaMKII increases during the DC injection, it is still unclear whether the activity of oCaMKII indeed helps the model learn or, in other words, what the exact role oCaMKII plays in priming. To investigate this, we perform a perturbation experiment to analyze the system in more detail. Specifically, we increase the initial activity of oCaMKII while conserving the total concentration of *cCaMKII* (neither open nor phosphorylated CaMKII) and oCaMKII, to see how the magnitude of the concentration of oCaMKII modulates the response of the whole system to the same stimulus.

In our set-up of the computational model, a single plateau potential (of the particular amplitude and duration that were used in the protocol) cannot induce the system to enter an LTP state. However, as shown in Fig.6B (left), the plateau potential combined with a perturbation that increases the initial concentration of oCaMKII by 4.8muM (about 12%) is enough to induce LTP, which confirms the critical role of oCaMKII in our model. Moreover, as also presented in Fig.6B (right), we can delay the onset of the plateau potential (for example, with a delay of 2.4 seconds), and LTP will still be induced – provided oCaMKII is initially increased by 100%. The required increase of oCaMKII and the duration of 2.4 seconds is consistent with BTSP, which suggests possible roles of both the priming of the activity of oCaMKII and the timescale of this activity in determining the timescale of BTSP.

The activity of oCaMKII is shown to play a vital role in priming, but it does not rule out the possibility of pCaMKII contributing to priming. Hence, we repeat the perturbation experiment for pCaMKII, and the results are shown in Fig.6C. It can be observed that an immediate increase of pCaMKII can also amplify the response of the system to the plateau potential and lead it to the LTP state, which is consistent with expectation. Besides, a relatively significant increase of pCaMKII paired with plateau potential 1.5 seconds later can also induce an LTP state. Although the effect decays faster than the previous controlled increase of oCaMKII, increasing pCaMKII can lead to an amplified response to stimuli. Also, the concentration of oCaMKII can increase during the decay of pCaMKII, as part of pCaMKII is transferred to oCaMKII. This increased oCaMKII can be present for a longer time, given that oCaMKII decays more slowly than pCaMKII.

## Discussion

We have developed and implemented a computational model of the biochemical reactions responsible for E-LTP/D for synapses of CA1 pyramidal neurons in the hippocampus. The model is a single compartment model of the post-synaptic density (PSD) of the spine head, containing well-mixed *Ca*^2+^ ions, CaM, CaMKII, and phosphatase. The biochemical reactions of the two interacting pathways [the kinase CaMKII pathway and a phenomenological phosphatase PP pathway] are described by a deterministic system of ordinary differential equations. The ion inflow (including *Ca*^2+^) is described by ion channels – NMDAR, CaV1, and Na channels. The dynamics is tri-stable, with three stable fixed points that represent the basal, LTP, and LTD states, together with the transient dynamics that describes the phosphorylation of CaMKII (that we identify with the formation of E-LTP), and the dephosphorylation of phosphatase (that we identify with the formation of E-LTD). As in *in-vitro* experiments, LTP/D can be induced by distinct signaling protocols, with STDP (spike timing dependent plasticity) induced by many repetitions of pre-post synaptic spike pairs and BTSP (behavioral time scale plasticity) induced by a few pre-synaptic spikes followed or preceded by a plateau potential.

Our model is the first (to our knowledge) to realize both STDP and BTSP in one model of the biochemistry in the PSD, with one identical set of parameter values. The model captures realistic responses to each signaling protocol (temporal profiles of the transient approaches to E-LTP, with proper timescales and Hebbian nature): for STDP, a causal (asymmetric) Hebbian temporal profile, with inter-pair time scales of tens of ms; for BTSP, a non-Hebbian temporal profile, with transient time scales of several seconds. Moreover, analysis of simulations identifies, within the model, differing primary mechanisms underlying STDP and BTSP: the *Ca*^2+^ flow sets the ms time scale and the causal Hebbian properties of STDP; the CaMKII activity sets the slower time scales and non-causal nature of BTSP. In addition, the model supports a priming mechanism that is induced by a small depolarization sufficient to open CaV1.3 channels for *Ca*^2+^ ion flow into the spine head. Once in the PSD, this *Ca*^2+^ opens the closed compact state of CaMKII, placing CaMKII in the ready for activation (phosphorylation) which triggers the induction of LTP, as well making it easier to realize BTSP. Thus, the model confirms expectations that NMDAR is responsible for the time scales of Hebbian plasticity of STDP, and that *Ca*^2+^ flow through CaV1 channels could enable a priming mechanism. In addition, the model clearly identifies the interaction of the timescales of the activation of CaMKII with those of the signaling protocols that result in STDP, BTSP, and priming.

Currently, it is not clear experimentally whether co-existing stable fixed points represent either early LTP or the fully phosphorylated state of active CaMKII. In the case of early LTP, the existence of bistable synapses is supported by experimental work showing an all-or-nothing transition of synaptic efficacy (41, 42); however, other experimental studies suggest synapses with a more graded degree of potentiation (43, 44). It also remains unclear experimentally whether stable E-LTP can be achieved by the biochemical activity of CaMKII and PP. Although many experiments and simulations observe and implement steady states of the CaMKII & PP system (45–48), newer findings show that the stable plasticity does not have to rely on the stability of the fully activated state of CaMKII (49–51). Here, we focus on modeling the induction of early LTP/D, over the first several minutes. Active CaMKII could be, and likely is, unstable over longer timescales (hours to days) due to succeeding reactions that are not included in our current model. We view the tri-stable fixed points in our model as representing states that are at best meta-stable, but with sufficient lifetimes to organize the transient dynamics of early LTP/D.

There are several differences of our model from previous models. *First*, to model the binding and disassociation of molecules and how they influence phosphorylation and dephosphorylation, we replace Michaelis-Menten schemes (quasi-steady-state) with quasi-equilibrium representations, which are a more fundamental description of the catalytic reactions. *Second*, our representation of the activity states of CaMKII includes the *compact state* of CaMKII, together with the transition between this compact state and an open but inactive state of CaMKII that is later activated. The presence of this compact state has been well established experimentally (16, 52, 53) and plays an important role in the model, where inducing a transition from compact to open increases the sensitivity of CaMKII to mild stimuli, and thus strengthens its role as an information integrator, on which we will elaborate below. *Third*, the model describes the kinase CaMKII pathway interacting with a phosphatase PP pathway, in which the bio-chemical reactions of the CaMKII pathway are described rather realistically (27), while the PP pathway is treated as a generic phosphatase pathway. Details of the particular phosphatases relevant for CA1 pyramidal neurons are still being studied experimentally, and seem to us somewhat open. For example, the commonly studied phosphatase inhibitor-1 (I-1) has been shown to have no effect on the PP1-mediated regulation of NMDAR-dependent synaptic plasticity in CA1 pyramidal neurons (54–56). Thus, we decided to represent the phosphatase pathway rather generically, adapting a phenomenological representation from Pi and Lisman (28). Our model of the two interacting pathways retains the tri-stability of the fully phenomenological model of Pi and Lisman – creating a straightforward representation of the synapse’s basal, LTP, and LTD states without ambiguity, and allowing a clear distinction between depression (transition from basal to LTD) and depotentiation (transition from LTP to basal).

It is widely known that CaMKII plays essential roles in synaptic plasticity, especially in early LTP & LTD. For example, experiments show that active CaMKII binds to the NMDAR and AMPAR channels, where it leads to a strengthening of the synapse (16, 17). In addition, there is experimental evidence that CaMKII can act as a frequency decoder in synaptic plasticity (57, 58). Recent experiments also suggest the critical involvement of CaMKII in BTSP (59). Our model highlights the potential importance of the seconds-long time scale of the transient activation of CaMKII. During this time, CaMKII can be partially activated through a weak stimulus, including pre-synaptic spikes and DC, making LTP easier to be induced by a more potent stimuli (for example, a plateau potential which can be induced by current injection or naturally occur, see (10)) which follows[Fig.6]. This transient activity can sustain for seconds and be further prolonged through phosphorylation of CaMKII, consistent with experimental findings (49, 50). The similarity between the timescale of CaMKII and the timescale of BTSP is intriguing: In the model’s description of BTSP, the activity of CaMKII successfully integrates information of pre-synaptic spikes and the plateau potential, spaced seconds apart. Moreover, again in the model, the time scales of active CaMKII play a vital role in STDP: With a seconds-long duration of activity, CaMKII is retained during the one-second intervals between the spike pairs (STDP utilizes a 1-Hz stimulus protocol) and accumulates within the large number of repetitions. In summary, the seconds-long time scale of the transient activation of CaMKII allows it to function as an information integrator, both sensitively and stably, making it suitable for different signaling patterns. In other words, the transient activation of CaMKII could operate as an *eligibility trace* for learning (39, 60, 61).

Our model also realizes a proposed theory of priming (26) in which CaMKII is mildly activated, making it easier for a strong stimulus to induce LTP. In the model, priming is activated by a small DC that selectively opens CaV1.3 channels, allowing for mild *Ca*^2+^ inflow which triggers the activation of CaMKII. The selectivity for CaV1.3 is thought to occur because, in the voltage range of the mild depolarization, CaV1.3 has a much steeper voltage dependence than CaV1.2 or NMDAR (26). With the model, we investigate the potential role of CaV1.3 channels in priming: First, we measure an enhanced calcium inflow through CaV1 channels (primarily CaV1.3) opened by the mild depolarization. This increases the baseline calcium concentration, which in turn enhances CaMKII activity. Within the model during priming, inflow through CaV1.3 channels is the primary contributor to the mild increase in calcium concentration, even though only a small portion (around 20%) of the CaV1 channels in the spine head is CaV1.3 (62, 63). We further highlight the role of CaV1.3 channels within the model by showing that the deactivation of CaV1.3 abolishes priming and obstructs learning [Fig.4, Fig.S8C]. Biologically, there are several possible ways that a CaV1 priming effect might be strengthened. For instance, the mild increase in baseline calcium might be further amplified by a calcium induced calcium release from the spine apparatus through RyR channels (64).

The model also shows how priming could play a role in behavioral plasticity (like BTSP). In the model, priming through DC injections extends the timescale of BTSP [Fig.S1B], consistent with experimental findings that increasing baseline membrane potential will increase the timescale of BTSP (39). Analyzing the model’s response [Fig.4] shows that priming increases the concentration of open CaMKII and slows its decay, which further prolongs CaMKII activity as an information integrator.

Within our model, a plateau potential induced plasticity protocol such as BTSP, awaken and stabilize place cells – effects which are aided by priming. In reality, this priming could arise from several possible sources – for example, a small DC injection or a transient ramp-like depolarization of place cells. Priming through ramp-like depolarization could explain the irreversible components in experiments of (24). Alternatively, a depolarizing current from other afferent neurons could function as a signal modulating the plasticity, providing a possible explanation for why a mouse shows different learning between familiar and novel settings (65). In any event, the model predicts that weak priming will lead to an LTP that can easily be reversed, and stronger priming will result in a more robust LTP.

In effect, priming alters the properties of the basal state, and hence the properties of the transition from the basal to the LTP state. This transition characterizes the properties of E-LTP – to study the induction of E-LTP (which occurs through the transition), it will be important to introduce stochasticity. Hence, we introduced a phenomenological noise term that perturbs the ion flow through the channels, in a manner similar to previous work (66–68). As shown in Results [Fig.2,Fig.3,Fig.5], in this stochastic setting, the properties of these transients are quite evident, with the transition probabilities between different states of CaMKII encoding how the synapse efficacy changes during different stimuli. In future work, this phenomenological noise will be replaced by a more detailed stochastic description of the biochemical reactions within the very small spine head, providing a more realistic representation of the stochastic dynamics. In this more realistic representation, stochasticity will arise from stochastic inflow through the ion channels, as well as from the small discrete number of ions and proteins present in the spine head.

Finally, our model can be generalized in several directions. In addition to discreteness effects with stochastic representations, the effects of the nonuniformity of the calcium concentration throughout the spine head must be captured. There is a much higher concentration near the mouth of the open channels, a spatial region of the spine head termed as the *calcium nanodomain* (69, 70). This calcium nanodomain will influence the local strength of calcium dynamics (71). Also, CICR (calcium induced calcium release) from the internal storage could play an important role in plasticity for neurons with spine apparatus (72). Describing the effects of such spatial inhomogeneities will require a spatially extended description of the spine head, including at least a multi-compartment representation approximating calcium diffusion. In addition, CaMKII and PP are known to be localized in the PSD (19, 73) and interact with ion channels (NMDAR, AMPAR, LTCC) through conformational changes there (25, 74–77). The conformational signaling and localization will also require a compartmental model distinguishing PSD and the rest of the spine head.

A better more realistic description of the LTD pathway is also needed. Given the diversity of experimental results, several pathways may be involved in the induction of LTD (20, 78); a more detailed model of calcium-inducing LTD must await further experimental studies.

Last but not least, an extension beyond a single synapse to multiple synapses and multiple neurons will be needed to model the emergence of place cells. For example, recent experimental work (39) unveils roles for multi-synapses/neurons in BTSP. Understanding the mechanisms underlying the roles of these multi-synapse/neurons will require the investigation of the communication among multiple synapses (the local depolarization, dendritic spikes, diffusion of CaMKII) and how this communication produces the global learning through coordination/modulation among multiple synapses.

## Materials and Methods

We utilize a single-compartmental, deterministic representation by a system of ordinary differential equations (ODEs) to model the biochemical reactions of early LTP/D at the post-synaptic density (PSD) region of a single synapse onto a post-synaptic dendritic spine. The synaptic events will depolarize the membrane and open the relevant ion channels, allowing calcium ions to flow into the spine head. The increasing calcium concentration will activate CaMKII and phosphatase, which will further activate themselves through autocatalytic reactions, while the two will inhibit each other through phosphorylation and dephosphorylation. This sequence of chemical reactions culminates in a strengthening (weakening) of the synaptic strength known as LTP (LTD). The general reaction scheme is illustrated in Fig.1A.

We describe the model in two parts: i) the dynamics of membrane potential and intracellular calcium concentration, including the dynamics of ion channels; ii) the biochemical reactions composed of two interacting pathways – one “UP” pathway that strengthens the synapses through a kinase called CaMKII, and one “DOWN” pathway that weakens the synapses through a phosphatase (phenomenologically described), combined with a “read-out” mechanism of LTP and LTD from the activity of CaMKII and phosphatase. A complete and detailed description of the model and its parameters is provided in the SI.

### Membrane potential and calcium signaling

To model the chemical system under different stimulus protocols, we need to investigate how the synaptic events induce calcium concentration changes in the spine head. To describe the dynamics of the membrane potential, we follow Graupner and Brunel (27) and embed the spine head in the whole cell.

### Membrane potential

We use a Hodgkin-Huxley formalism model to describe the membrane potential *V* with the following equation:

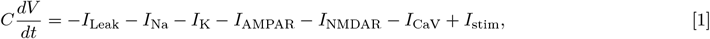

where *C* is the membrane capacitance, and *I*_*x*_ are the ion currents from different sources. . (1) describes the voltage V as an iso-potential for the whole cell, including both the soma and the dendrite. Accordingly, capacitance, leakage conductance, and the conductance of sodium and potassium channels are all selected with respect to a surface area of 10000*μm*^2^ (roughly the surface area of a pyramidal neuron), ensuring the generation of a realistic action potential. Moreover, the AMPAR channel conductance is scaled up by tenfold compared to its typical value on one single spine head, resulting the generation of an EPSP around 1mV—a value consistent with experimental observations at the soma. The NMDAR and calcium channels represent ion flow into the spine head. We keep these two conductances normalized to a single spine head (no scaling up), to produce reasonable calcium dynamics within the spine head. In the single compartment iso-potential model, there is no distinction between the voltage at the spine head and that at the soma; thus, in our model the NMDAR and CaV synaptic input at the spine head only contribute negligibly to EPSPs at the spine head.

### Voltage-dependent calcium channels

Voltage-gated calcium channels (CaV1.2, CaV1.3) potentially play an important role in the mechanisms underlying LTP/LTD. A model beyond a Hodgkin-Huxley representation is required as these channel types show calcium-induced inactivation. Thus we take a Markovian model of CaV channels in the rabbit heart cell (79), and adapt it to CaV1.2,CaV1.3 channels in the spine setting (80, 81).

### Calcium signaling in a dendritic Spine

Calcium concentration change in the PSD is modeled in a single compartment uniformly in space, similar to (27). Two main sources of calcium inflow into the PSD are calcium influx through NMDAR and voltage-gated calcium channels. The calcium concentration follows the differential equation:

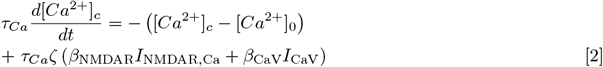

where *ζ* is the factor converting the calcium current to calcium concentration change per unit time and *β*_NMDAR_, *β*_CaV_ are the fast buffering factor of calcium influx through NMDAR and CaV channels, respectively. The extrusion, diffusion, and slow buffering of calcium are not modeled explicitly, but described by a leak term with single exponential decay to the resting calcium concentration [*Ca*^2+^]_0_ with the time constant *τ*_*Ca*_ equal to 12 ms.

### Calcium trace with noise

The stochastic simulations are performed by adding noise to the calcium dynamic during the synaptic events, through the stochastic opening and closing of ion channels. To represent this stochastic effect, we include two sources of noise following (27): i) random maximum conductance of NMDAR at each presynaptic spike and ii) random maximum conductance of the CaV channels at each postsynaptic spike and plateau potential. During each synaptic event, the channel conductance is drawn from a Gaussian distribution with a given mean and variance, chosen so that the stochastic system can reproduce smooth results, consistent with experiments.

### Experimental stimulus protocols

A prescription of presynaptic and postsynaptic spikes and postsynaptic plateau potentials are required to simulate the system under different experimental protocols. Instead of modeling spiking as in vivo, we initiate the system with prescribed presynaptic and postsynaptic spike trains (and plateau potentials).

In our model, we do not detail the presynaptic spiking, the release of neurotransmitters at the presynaptic side, the diffusion of transmitters into the synaptic cleft, or their capture by glutamate receptors. Consequently, synaptic failure is not considered. Instead, we only describe the binding of neurotransmitters to AMPARs and NMDARs. After each presynaptic spike, each AMPAR and NMDAR will bind to one neurotransmitter immediately, initiating channel activation (shown in SI). For the postsynaptic spikes, we generate an artificial action potential by applying a 1ms pulse current of 3000 pA, which causes a spike approximately 2ms after the current injection. Similarly, the postsynaptic plateau potential is induced by a sustained current of 4000 pA (3000 pA for the stochastic scenarios), typically lasting for 300 ms.

### Biochemical reactions in a single chamber

The model of biochemical reactions is based on a model of early LTP/D (27) as a bistable system of ordinary differential equations (ODEs), with both UP and DOWN pathways. The model describes the calcium-induced activation of both the kinase (CaMKII) and the phosphatase and the mutual inhibition between the two, with detailed reaction scheme illustrated in Fig.1 and Fig.S4. Starting from the model in (27), we replace several components in the UP kinase pathway with a more realistic and precise description (adding a compact state of CaMKII (52) and using quasi-equilibrium instead of a Michaelis-Menten scheme). We remove the traditional I1-PP1 signaling for the DOWN phosphatase pathway (as it is reported not to be observed in CA1 pyramidal neurons (54, 82, 83)) and substitute with a phenomenological representation (28). This resulting model is tri-stable with three coexisting stable steady states, directly related to the synapse’s basal, LTP & LTD states.

### Binding and disassociation in fast equilibrium

The inflowing calcium will bind with Calmodulin (CaM), and the calcium-CaM complex can further bind to CaMKII and trigger its activation. At the same time, the phosphorylation of CaMKII requires the binding of ATP and the dephosphorylation of CaMKII requires the binding of phosphatase. We introduce a *fast-equilibrium* assumption to describe the binding and disassociation between molecules. Specifically, we assume that for the binding of molecular X with Y,

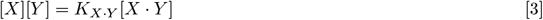

holds for all time, where [*X*], [*Y*], [*X · Y*] are the concentration of free *X*, free *Y* and *X · Y* complex, respectively and *K*_*X*_*·*_*Y*_ is a dissociation constant independent of time.

Calmodulin (CaM) is a calcium-binding messenger protein with two lobes (the N- and C-domains), and each lobe can bind to at most two calcium ions. The resulting calcium-CaM complex can further bind to CaMKII and trigger its activation. For simplicity, we only consider the number of calcium ions bound to Calmodulin without distinguishing between states with different binding sites but the same number of calcium ions bound. This simplification leads to five states of CaM (from zero to four ions bound), similarly to (27). Fast-equilibrium is used to model the binding and disassociation of calcium ions, leading to four algebraic equations (shown in supporting information).

CaMKII is a dodecameric holoenzyme composed of two stacked hexameric rings formed by six functionally coupled subunits (84). Each subunit provides multiple binding sites for different molecules, and the binding can promote the phosphorylation or dephosphorylation of each subunit, as illustrated in Fig.S4A. In our model, we also assume that the binding and dissociation of molecules (Calcium-CaM complex, phosphatase, ATP, ADP) with CaMKII is in fast-equilibrium, similar to calcium binding to Calmodulin (detailed equations presented in supporting information). The fast-equilibrium assumption, combined with the description of how the binding catalyzes reactions (discussed in the next section), provides an alternative model to describe the catalyzation of phosphorylation and dephosphorylation of CaMKII subunits instead of the Michaelis-Menten scheme used in (27).

### Phosphorylation and dephosphorylation of CaMKII

The binding of the calcium-CaM complex will initiate the phosphorylation at threonine-286 (Thr-286) in the regulatory domain of CaMKII, which disrupts the binding of the regulatory domain to the kinase domain (84). After being phosphorylated, CaMKII subunits can stimulate the phosphorylation of their neighboring subunits, prolonging the activation after the calcium stimulus (85). In our model, CaMKII activity serves as an indicator of potentiation. It is treated in some detail, but with some aspects not considered for simplicity, such as different isoforms of CaMKII subunits or the phosphorylation at Thr-305 and 306.

In modeling the phosphorylation and dephosphorylation of CaMKII, we closely follow Graupner and Brunel (27). Since little is known about how the two rings of the CaMKII interact and how that affects phosphorylation (86), we suppose the two are independent of each other and model only one six-subunit ring. Each CaMKII kinase is treated as a six-subunit ring in which each subunit can be phosphorylated or unphosphorylated independently. This results in a total of 14 states under rotation invariance, as shown in Fig.S4C. The transition between these different states is through the phosphorylation and dephosphorylation of subunits, which can be divided into two categories (illustrated in Fig.S4,A and B) : i) the spontaneous phosphorylation and dephosphorylation of one subunit and ii) the neighboring phosphorylation of CaMKII (and its reverse reaction). We present the detailed equations and parameters in the supporting information.

### Docking and undocking of CaMKII

The presence of the compact state of CaMKII has been well established experimentally (16, 52, 53, 87). Compact CaMKII can be bound to actin filaments in the spine head and dendritic shaft, where the compact state’s release from the actin is triggered by calcium-calmodulin binding (25). In our model, we include this compact state as an additional state of CaMKII. Compact CaMKII can become open (“undocking”) with a calcium-dependent rate, and the open CaMKII can return to the compact state (“docking”) when it has no phosphorylated subunits and is not bound to calmodulin or phosphatase [Fig.S4D].

The transition between the compact state and open state is given by

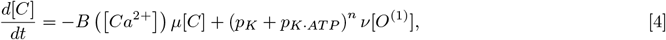

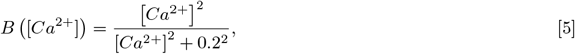

where [*C*] and [*O*^(1)^] are the concentration of the CaMKII in the compact state and the open state with no phosphorylated subunits, respectively, and *p*_*K*_ + *p*_*K AT P*_ is the probability that one unphosphorylated subunit is not bound to calmodulin or phosphatase.

### Phenomenological phosphatase dynamics

Details of the particular phosphatases relevant for CA1 pyramidal neurons are still being studied experimentally (20), and seems to us somewhat open. Thus, we decided to represent the phosphatase pathway rather generically, adapting a phenomenological representation from Pi and Lisman (28):

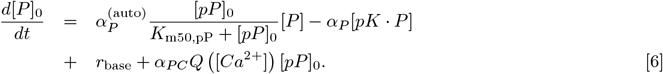

with [*P*]_0_ + [*pP*]_0_ = [*P*]_*tot*_, [*P*] + [*pK · P*] = [*P*]_0_ and

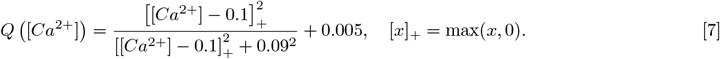

[*P*]_0_ and [*pP*]_0_ are the concentrations of active (dephosphorylated) and inactive (phosphorylated) phosphatase, respectively and *r*_base_ is the baseline dephosphorylation activity rate. *α*_*P C*_*Q* [*Ca*^2+^] describes the calcium dependent rate of dephosphorylation with a lower activation threshold (compared with CaMKII).

### LTP, LTD, and synaptic efficacy

In the model, LTP is represented by the dominance of the concentration of active CaMKII over active PP, and LTD by the dominance of the concentration of active PP over CaMKII. As an indicator of the resulting synaptic efficacy, we adopt a heuristic ‘read-out’ representation [taken from (28)] of the ratio of the number of AMPA receptors on the spine head before and after induction:

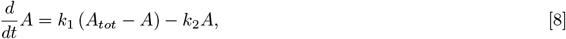

where *A* is the number of AMPA receptors on the membrane, and *A*_*tot*_ is the number of total available AMPA receptors. In this heuristic representation of AMPA trafficking, *k*_1_ (*k*_2_) is the rate AMPA is inserted in (removed from) the membrane, each represented by a linear function:

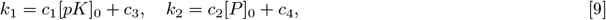

where *c*_1_, *c*_2_ are the scaling factors, and *c*_3_ and *c*_4_ are the insertion and removal rates independent of CaMKII and phosphatase activities. We emphasize that Eq. (8) operates on a much slower time scale than the activation of CaMKII and PP; thus, Eq. (8) does not couple back to the dynamics of the CaMKII and PP, but is just a read-out from the concentrations of active CaMKII and PP, which provides an indication of synaptic efficacy.

## ACKNOWLEDGMENTS

We gratefully acknowledge Richard Tsien for discussions generally about hippocampal LTP/D at the molecular-cellular level, and in particlar about a possible role of CaV channels in a priming mechanism for LTP.

