## Supplementary for "A Biochemical Description of Postsynaptic Plasticity – with Timescales Ranging from Milliseconds to Seconds"

#### This PDF file includes:

Supplementary text
Figs. S1 to S9
Tables S1 to S2
SI References

### Supporting Information Text

**A Heuristic Intuition about the Model's Mechanisms Underlying several Aspects of BTSP.** In this section, we heuristically consider the chemical system we model as it undergoes transitions between the "basal" and "LTP" states under BTSP stimulation, seeking intuition about i) the effect of steady depolarization (DC injection) and ii) the origin of an observed asymmetry in the plots of synaptic efficacy vs $\Delta t$ – that is, an asymmetry with respect to the sign of $\Delta t$, the temporal delay between the pair of stimuli of the BTSP protocol (Fig. 3D, (1)).

The BTSP protocol consists in the composition of two stimuli (a presynaptic spike train and a postsynaptic plateau potential), separated in time by $\Delta t$. The sign of $\Delta t$ determines whether the presynaptic spike train precedes or follows the postsynaptic plateau. Thus, the "one-shot" BTSP stimulus is the composition (Stimulus 1 + $\Delta t$ delay + Stimulus 2), where Stimulus 1 can be either pre-synaptic spike train or the post-synaptic plateau, and Stimulus 2 the opposite choice. A steady depolarization is also injected by a DC that is completely independent of Stimulus 1 and Stimulus 2.

We introduce an abstract and heuristic notion of an "activation state" of the system, where large values of this activation state indicate the dominance of active kinase over phosphatase, yielding LTP; while moderate values indicate a balance between kinase and phosphate, yielding the basal state. As the activation state crosses a threshold value (**Th**), the phase point of the chemical system will transition from the basin of attraction of the basal state to the basin of attraction of LTP. In the BTSP setting, the activation state depends upon the two stimuli, the steady depolarization by DC, and time delay between the two stimuli.

Denoting **the activation state of the chemical system after Stimulus 1 as**

$$F(S_1, DC),$$

we further assume **activation state after time interval ($\Delta t$)**

$$F(S_1, DC, \Delta t)$$

can be expressed as

$$F(S_1, DC, \Delta t) = F(S_1, DC)\ r(\Delta t, DC),$$

where $r(\Delta t, DC)$ describes the decay of the activation state, depending solely on the $\Delta t$ and $DC$.

We now consider the activation state immediately after the second stimulus $S_2$, denoted as

$$F\left[S_2, DC \mid F\left(S_1, DC, \Delta t\right)\right].$$

This activation state must exceed the threshold **Th** for LTP; that is, in order for the second stimulus $S_2$ to place the phase point of the system into the basin of attraction of LTP state,

$$F\left[S_2, DC \mid F\left(S_1, DC, \Delta t\right)\right] \geq \mathbf{Th}. \quad [1]$$

This condition can be transferred to a condition on the activation state just before the second stimulus $S_2$, by noting there must exist a threshold that is a function of the magnitude of Stimulus 2 (and $DC$)

$$th(S_2, DC)$$

such that

$$F\left[S_2, DC \mid F_1\right] \geq Th, \quad \forall F_1 \geq th(S_2, DC). \quad [2]$$

**The system can only transition to "LTP" state if its activation state**

$$F(S_1, DC, \Delta t) = F(S_1, DC)\ r(\Delta t, DC)$$

**crosses the threshold at the onset of Stimulus 2**; that is,

$$F(S_1, DC)\ r(\Delta t, DC) \geq th(S_2, DC). \quad \textbf{(LTP condition)} \quad [3]$$

***Effect of direct current.*** The heuristic description provides us the opportunity to investigate the effect of DC on LTP induction. First, we **approximate the decay of the activation state after first stimulus as an exponential decay**:

$$r(\Delta t, DC) = \exp(-|\Delta t|/\tau(DC)), \quad [4]$$

with time constant $\tau(DC)$. The "LTP condition" Eq. (3) can then be expressed as

$$F(S_1, DC)\exp(-|\Delta t|/\tau(DC)) \geq th(S_2, DC), \quad [5]$$

which leads to an upper bound for $\Delta t$, the time difference between the two stimuli beyond which there is no LTP:

$$|\Delta t| \leq \tau(DC) \cdot \log\left(\frac{F(S_1, DC)}{th(S_2, DC)}\right). \quad \textbf{(Upper threshold of timing difference)} \quad [6]$$

Eq. (6) describes how the timescale of BTSP (upper threshold of $\Delta t$) depends on $\tau(DC)$, and upon the magnitude of both stimuli (through $F(S_1, DC), th(S_2, DC)$). **With increasing** $DC$, it is reasonable to expect the following:

1. **stronger activity at the end of Stimulus 1**, so larger $F(S_1, DC)$,
2. **lower threshold required at the beginning of Stimulus 2**, so smaller $th(S_2, DC)$,
3. **slower decay of the activity**, so larger time constant $\tau(DC)$ (likely a secondary effect compared to the aforementioned two factors).

Combining three statements, we conclude that $\Delta t$ **is a monotonic increasing function of** $DC$. This implies that increasing the DC will extend the timescale of BTSP, as observed in the numerical experiments (Fig.S1(B)).

***The asymmetric timescale.*** We turn to investigate the observed asymmetry of timescales between "pre-post" and "post-pre" in BTSP. First, we assume that **the activation state is a linear response to stimulus** (and DC), which leads to

$$F(S_1, DC) = kS_1 + \gamma DC, \quad F\left[S_2, DC \mid F_1\right] = kS_2 + \gamma DC + F_1, \tag{7}$$

and (based on the definition of $th(S_2, DC)$, Eq. (2))

$$th(S_2, DC) = \mathbf{Th} - (kS_2 + \gamma DC)\,. \tag{8}$$

Combined with the definition of upper bound of $\Delta t$ (Eq. (6)), we have the new formula of upper bound as:

$$\Delta t \leq \tau(DC) \cdot \log\left(\frac{F(S_1, DC)}{th(S_2, DC)}\right) = \tau(DC) \cdot \log\left(\frac{kS_1 + \gamma DC}{\mathbf{Th} - (kS_2 + \gamma DC)}\right). \tag{9}$$

First, we propose a reasonable assumption: **plateau potential can induce a stronger response than the pre-synaptic spike trains**,

$$S_{pre} < S_{post}. \tag{10}$$

We further assume **the system can reach "LTP" state when there is no delay time between the pre-synaptic and post-synaptic stimuli**, which leads to

$$F(S_{pre}, DC) \geq th(S_{post}, DC), \tag{11}$$

which is equivalent to

$$(kS_{pre} + \gamma DC) + (kS_{post} + \gamma DC) \geq \mathbf{Th}. \tag{12}$$

Now we turn to consider the following difference of two products; we have that

$$\begin{aligned}
& (kS_{pre} + \gamma DC) \cdot (\mathbf{Th} - (kS_{pre} + \gamma DC)) - (kS_{post} + \gamma DC) \cdot (\mathbf{Th} - (kS_{post} + \gamma DC)) \\
= \; & (kS_{post} + \gamma DC)^2 - (kS_{pre} + \gamma DC) - \mathbf{Th} \cdot ((kS_{post} + \gamma DC) - (kS_{pre} + \gamma DC)) \\
= \; & ((kS_{post} + \gamma DC) + (kS_{pre} + \gamma DC))\,((kS_{post} + \gamma DC) - (kS_{pre} + \gamma DC)) \\
- \; & \mathbf{Th} \cdot ((kS_{post} + \gamma DC) - (kS_{pre} + \gamma DC)) \\
= \; & [(kS_{post} + \gamma DC) + (kS_{pre} + \gamma DC) - \mathbf{Th}] \cdot k\,(S_{post} - S_{pre})\,.
\end{aligned} \tag{13}$$

Based on Eq. (10) and Eq. (12), we have that

$$[(kS_{post} + \gamma DC) + (kS_{pre} + \gamma DC) - \mathbf{Th}] \cdot k\,(S_{post} - S_{pre}) \geq 0, \tag{14}$$

which is equivalent

$$\frac{kS_{pre} + \gamma DC}{\mathbf{Th} - (kS_{post} + \gamma DC)} \geq \frac{kS_{post} + \gamma DC}{\mathbf{Th} - (kS_{pre} + \gamma DC)}. \tag{15}$$

It further gives out

$$\tau(DC) \cdot \log\left(\frac{kS_{pre} + \gamma DC}{\mathbf{Th} - (kS_{post} + \gamma DC)}\right) \geq \tau(DC) \cdot \log\left(\frac{kS_{post} + \gamma DC}{\mathbf{Th} - (kS_{pre} + \gamma DC)}\right). \tag{16}$$

The left term of Eq. (16) is the threshold of $\Delta t$ for $S_1 = S_{pre}, S_2 = S_{post}$ (pre-synaptic spike trains followed by plateau potential) and the right term is the threshold of $\Delta t$ for $S_1 = S_{post}, S_2 = S_{pre}$ (plateau potential followed by pre-synaptic spike trains). Eq. (16) indicates that the **timescale for "pre-post" stimulus is larger than "post-pre" stimulus**, providing an explanation for the **asymmetric timescale** in BTSP.

The analysis above suggests that the asymmetry of BTSP is inherited from the asymmetry of the "pre" and "post" stimulus' strength, leading to a prediction that varying the stimulus's strength can tune the asymmetry's strength. We have validated this prediction in our model, and the results are shown in Fig.S2. When fixing the post-synaptic plateau potential (the stronger stimulus) and increasing the strength of the pre-synaptic spike trains (with more spikes), we observe a more symmetric BTSP curve; vice versa; if we fix the pre-synaptic spike trains (the weaker stimulus) and increase the strength of the plateau potential (with a larger current injection), a more asymmetric BTSP curve is presented.

**Phosphorylation states of CaMKII.** In the main text, we have illustrated that the activity of the open and phosphorylated CaMKII have timescales different by three times (Fig. 6A). However, it is worth noting that CaMKII is a multi-subunit protein with many activation states (15 states within the model). It would be informative to track the dynamics of all the different states of the CaMKII. Here we group the states of CaMKII by the number of phosphorylated subunits and investigate the evolution of the CaMKII with different phosphorylation levels in several scenarios.

***Decay and timescales.*** We start estimating the timescales based on the same perturbation experiment in Fig.6A while focusing on the timescale of CaMKII with different phosphorylation levels instead of the phosphorylated and dephosphorylated subunits. The results are shown in Fig.S3A, and we can see that the activities decay faster with the higher phosphorylation level, consistent with previous observations that open CaMKII subunits have a much longer timescale compared with the phosphorylated ones. This finding also verifies that the transient, mild activation of CaMKII induced by subthreshold depolarization could retain for a longer time. It indicates the vital role of transient activation of CaMKII after subthreshold depolarization – an implementation of "priming". Besides, the diverse timescales enable CaMKII to be a flexible information integrator that can capture various stimulus patterns.

***Induction of LTP.*** We now turn to investigate the dynamics of CaMKII during the induction of LTP, which also shows how the different CaMKII states will provide us with more information about the dynamics. It is observed in our model (as shown in Fig.6, B and C) that a plateau of phosphorylated CaMKII activity exists after a relatively weak stimulus (for example, the combination of plateau potential with perturbation of the initial state in Fig.6, B and C), followed by a dramatically increasing phosphorylation.

The dynamics of CaMKII with different phosphorylation levels are shown in Fig.S3B, where the induction of LTP can be separated into three phases. During phase I, weakly phosphorylated CaMKII (with $\leq 3$ phosphorylated subunits) builds up slowly through weak calcium-induced activation and weak neighboring phosphorylation, corresponding to the plateau between 0 and 20 seconds in Fig.6B. However, a significant increase of strongly phosphorylated CaMKII (with $> 3$ phosphorylated subunits) is observed in phase II, indicating a strong activation of CaMKII through intense neighboring phosphorylation. Later in phase III, the activity of CaMKII gets stabilized and approaches the steady state.

The above results suggest how CaMKII could facilitate the multi-subunit structure to process different stimuli and initiate the strong activation, which can not be fully described merely with the dynamics of phosphorylated and open CaMKII. Moreover, future experimental observations with finer spatiotemporal resolutions (focused on CaMKII with different phosphorylation levels) could greatly deepen our understanding of the role of CaMKII in information integration and the induction of LTP.

### Methods and Model

Our model is a single compartment representation of the post synaptic density (PSD) region of the spine head. More specifically, we model the bio-chemical reactions in the PSD that are triggered by $Ca^{2+}$ ions flowing through channels on the membrane – channel flows that are induced by time dependent changes in the pre and post synaptic membrane potential. This sequence of chemical reactions culminates in a strengthening (weaking) of the synaptic strength known as LTP (LTD). We describe the model in three parts: i) the dynamics of the ion channels, including the dynamics of the membrane potential; ii) the dynamics of the bio-chemical reactions within the PSD, including the activation of the calcium/calmodulin-dependent protein kinase II (CaMKII) and the phosphatase (phenomologically described); and iii) a "read-out" of LTP and LTD from the concentrations of active CaMKII and phosphatase. First, we describe the membrane potential and channel dynamics.

**Hodgkin-Huxley Membrane and its Channels.** To model the chemical system under different stimulus protocols, we need to know how the membrane potential (across the membrane of the spine head) changes during the stimulus and how it induces calcium concentration changes in the PSD. This part of the model is standard, similar to that used in Graupner, and Brunel (2) with some modifications of the parameters of the ion channels.

***Membrane potential.*** The membrane potential $V$ obeys the following Hodgkin-Huxley formalism:

$$C_m \frac{dV}{dt} = -I_{\text{Leak}} - I_{\text{Na}} - I_{\text{K}} - I_{\text{AMPAR}} - I_{\text{NMDAR}} - I_{\text{CaV}} + I_{\text{stim}} \qquad [17]$$

where $C_m$ is the membrane capacitance, and $I_x$ are the ion currents from different sources.

***Ion currents.*** Following (2), we take the parameters of the membrane ion currents $(I_{\text{Leak}}, I_{\text{Na}}, I_{\text{K}})$ from (3) and the synaptic ion currents $(I_{\text{AMPAR}}, I_{\text{NMDAR}})$ from (4). We use a Markovian model for voltage gated Calcium channels $(I_{\text{CaV}})$, modified from (5), instead of a Hodgkin-Huxley type model in (2).

**Leak Current** The leak current is modeled as

$$I_{\text{Leak}} = g_{\text{Leak}}(V - E_{\text{Leak}}), \qquad [18]$$

where $g_{\text{Leak}}$ is the leak conductance and $E_{\text{Leak}}$ is the leaking potential.

**Sodium current** The sodium current is modeled with a Hodgkin-Huxley sodium channel with two gating variables. In a Hodgkin-Huxley model, any gating variable $x$ will obey the first-order dynamics with voltage-dependent rates:

$$\frac{dx}{dt} = \left(x_\infty(V) - x\right)/\tau_x(V). \qquad [19]$$

The sodium current is given by

$$I_{\mathrm{Na}} = g_{\mathrm{Na}} m_{\mathrm{Na}}^3 h_{\mathrm{Na}} (V - E_{\mathrm{Na}}), \qquad [20]$$

while the gating variables satisfy that

$$m_{\infty \mathrm{Na}}(V) = \frac{1}{1 + e^{-(V+36.5)/5}}, \quad \tau_{m_{\mathrm{Na}}} = 0.1\ ms, \qquad [21]$$

$$h_{\infty \mathrm{Na}}(V) = \frac{1}{1 + e^{(V+44.1)/7}}, \quad \tau_{h_{\mathrm{Na}}} = \left(\frac{3.5}{e^{(V+35)/4} + e^{-(V+35)/25}} + 1\right)\ ms. \qquad [22]$$

**Delayed-rectifier potassium current** The model of potassium current is a Hodgkin-Huxley model similar to sodium current with only one gating variable. It is described by

$$I_{\mathrm{K}} = g_{\mathrm{K}} n_{\mathrm{K}}^4 (V - E_{\mathrm{K}}), \qquad [23]$$

with

$$n_{\infty \mathrm{K}}(V) = \frac{1}{1 + e^{-(V+30)/25}}, \quad \tau_{n_{\mathrm{K}}} = \left(\frac{2.5}{e^{(V+30)/40} + e^{-(V+30)/50}} + 0.01\right)\ ms. \qquad [24]$$

**AMPAR current** Spiking of the presynaptic neuron will lead to the release of neurotransmitters, which further open the AMPA receptor and NMDA receptor on the postsynaptic membrane. The AMPA receptor-mediated current is described by

$$I_{\mathrm{AMPAR}} = g_{\mathrm{AMPAR}} s_{\mathrm{AMPAR}} \left(V - E_{\mathrm{AMPAR}}\right), \qquad [25]$$

$$\dot{s}_{\mathrm{AMPAR}} = -s_{\mathrm{AMPAR}}/\tau_{\mathrm{AMPAR}} + \alpha_s x_{\mathrm{AMPAR}} \left(1 - s_{\mathrm{AMPAR}}\right), \qquad [26]$$

$$\dot{x}_{\mathrm{AMPAR}} = -x_{\mathrm{AMPAR}}/\tau'_{\mathrm{AMPAR}} + \alpha_x \sum \delta(t - t_k), \qquad [27]$$

while $\{t_k\}$ is the sequence of presynaptic spiking time and $\delta$ is the Dirac function.

**NMDAR current** The NMDA receptor-mediated current is given by

$$I_{\mathrm{NMDAR}} = g_{\mathrm{NMDAR}} s_{\mathrm{NMDAR}} B(V) \left(V - E_{\mathrm{NMDAR}}\right), \qquad [28]$$

while the magnesium block is modeled as

$$B(V) = \frac{1}{1 + e^{-0.062V} \frac{[Mg^{2+}]}{3.57}}. \qquad [29]$$

The gating variable $s_{\mathrm{NMDAR}}$ satisfies the same type of dynamics as $s_{\mathrm{AMPAR}}$, but with different time constants $\tau_{\mathrm{NMDAR}}$ and $\tau'_{\mathrm{NMDAR}}$.

**Voltage-dependent calcium current** Voltage-gated calcium channels (CaV1.2, CaV1.3) potentially play an important role in the mechanisms underlying LTP/LTD. As these channel types have calcium-induced inactivation, a model beyond a Hodgkin-Huxley representation is required. The same subtypes of voltage-gated channels arise in cardiac cells, where they are well measured and modeled (6). We take a Markovian model of CaV channels in the rabbit heart cell (5), and adapt it to CaV1.2,CaV1.3) channels in the spine setting. Thus, the calcium current is given by

$$I_{\mathrm{CaV}} = g_{\mathrm{CaV}} P_o i_{\mathrm{Ca}}, \quad i_{\mathrm{Ca}} = \frac{4 P_{\mathrm{Ca}} V F^2}{RT} \frac{c_s e^{2a} - 0.341 \left[Ca^{2+}\right]_o}{e^{2a-1}}, \quad a = VF/RT, \qquad [30]$$

where $P_o$ is the open probability of the CaV channels and $c_s$ is the cytoplasmic calcium concentration satisfying Eq. (38).

The open probability $P_o$ is modeled with an ODE Markovian system modified, with some transition rates are adjusted to fit the experimental data in neuron cells (7, 8):

$$\begin{aligned}
\frac{dC_2}{dt} &= \beta C_1 + k_5 I_{2Ca} + k_5' I_{2Ba} - \left(k_6 + k_6' + \alpha\right) C_2, \\
\frac{dC_1}{dt} &= \alpha C_2 + k_2 I_{1Ca} + k_2' I_{1Ba} + r_2 P_o - \left(r_1 + \beta + k_1 + k_1'\right) C_1, \\
\frac{dI_{1Ca}}{dt} &= k_1 C_1 + k_4 I_{2Ca} + s_1 P_o - (k_2 + k_3 + s_2) I_{1Ca}, \\
\frac{dI_{2Ca}}{dt} &= k_3 I_{1Ca} + k_6 C_2 - (k_4 + k_5) I_{2Ca}, \\
\frac{dI_{1Ba}}{dt} &= k_1' C_1 + k_4' I_{2Ba} + s_1' P_o - (k_2' + k_3' + s_2') I_{1Ba}, \\
\frac{dI_{2Ba}}{dt} &= k_3' I_{1Ba} + k_6' C_2 - (k_4' + k_5') I_{2Ba}, \\
\frac{dc_p}{dt} &= \tilde{g}_{Ca} P_o i_{Ca} - \frac{c_p - [Ca^{2+}]_c}{\tau_s} \qquad [31]
\end{aligned}$$

where $c_p$ is the calcium concentration at the mouth of CaV channels and $P_o$ is the open probability of the single channels described as

$$P_o = 1 - (C_1 + C_2 + I_{1Ca} + I_{2Ca} + I_{1Ba} + I_{2Ba}), \quad [32]$$

The transition rates between Markovian states are given by

$$\alpha = p_o^{\infty}/\tau_{po},\ \beta = (1 - p_o^{\infty})/\tau_{po},\ p_o^{\infty} = \frac{1}{1 + e^{-(V+\Delta V)/8}}, \quad [33]$$

$$\begin{aligned} s_1 &= 0.02 f(c_p), \\ k_1 &= 0.03 f(c_p), \\ s_2 &= s_1 (k_2/k_1)(r_1/r_2), \\ s_2' &= s_1' \left(k_2'/k_1'\right)(r_1/r_2), \\ f(c_p) &= \frac{1}{1 + (\tilde{c}_p/c_p)^3}, \end{aligned} \quad [34]$$

$$\begin{aligned} k_3 &= \frac{e^{-(V+\Delta V+40)/3}}{3\left(1 + e^{-(V+\Delta V+40)/3}\right)}, \\ k_3' &= k_3, \\ k_4 &= k_3 (\alpha/\beta)(k_1/k_2)(k_5/k_6), \\ k_4' &= k_3' (\alpha/\beta)\left(k_1'/k_2'\right)\left(k_5'/k_6'\right), \end{aligned} \quad [35]$$

$$\begin{aligned} k_5 &= (1 - P_s)/\tau_{Ca}, \\ k_6 &= f(c_p) P_s/\tau_{Ca}, \\ k_5' &= (1 - P_s)/\tau_{Ba}, \\ k_6' &= P_s/\tau_{Ba}, \\ \tau_{Ca} &= (R(V + \Delta V) - T_{Ca}) P_r + T_{Ca}, \\ \tau_{Ba} &= (R(V + \Delta V) - T_{Ba}) P_r + T_{Ba}, \\ T_{Ca} &= \frac{78.0329 + 0.1\left(1 + (c_p/\bar{c}_p)\right)^4}{1 + (c_p/\bar{c}_p)^4}, \\ R(V) &= 10 + 4954 e^{V/15.6}, \\ P_r &= 1 - \frac{1}{1 + e^{-(V+\Delta V+40)/4}}, \\ P_s &= \frac{1}{1 + e^{-(V+\Delta V+40)/11.32}}. \end{aligned} \quad [36]$$

where $\Delta V$ is a constant that modifies the voltage dependency for different subtypes of CaV channels. It is noteworthy that there are two types of CaV channels present in the dendritic spine: CaV1.2 and 1.3. We include both subtypes and set $\Delta V = 0mV$ for CaV1.2 and $\Delta V = 30mV$ for CaV1.3 channels. With these choices, the I-V curves are consistent with experimental observations (7, 8). The total calcium current through CaV channels is a sum of the two sources:

$$I_{\mathrm{CaV}} = I_{\mathrm{CaV1.2}} + I_{\mathrm{CaV1.3}} \quad [37]$$

with each satisfying Eq. (30).

***Experimental stimulus protocols – presynaptic and postsynaptic membrane voltage profiles..*** A prescription of presynaptic and postsynaptic spikes and postsynaptic plateau potentials are required to simulate the system under different experimental protocols. Instead of modeling how the spike events in vivo, we initiate the system with pre-described presynaptic and postsynaptic spike trains (and plateau potentials).

The presynaptic spike trains are presented to the system through the release timing of neurotransmitters, which is encoded in Eq. (27) and the unshown equation of $x_{\mathrm{NMDAR}}$. For the postsynaptic spikes, we generate an artificial action potential by applying a 1ms pulse current of 3000 pA, while the spike occurs approximately 2ms after the current injection. Similarly, the postsynaptic plateau potential is induced by a sustained current of 4000 pA (3000 pA for the stochastic scenarios), typically lasting for 300 ms.

**Calcium Signaling in a Dendritic Spine.** Calcium concentration change in a dendritic spine (PSD) is modeled in a single compartment uniformly in space, similar to (2). Two main sources of calcium inflow into the PSD are calcium influx from NMDAR and voltage-gated calcium channels.

***Calcium dynamics.*** The calcium concentration follows the differential equation:

$$\tau_{Ca}\frac{d[Ca^{2+}]_c}{dt} = -\left([Ca^{2+}]_c - [Ca^{2+}]_0\right) + \tau_{Ca}\zeta\left(\beta_{\text{NMDAR}} I_{\text{NMDAR,Ca}} + \beta_{\text{CaV}} I_{\text{CaV}}\right) \quad [38]$$

where $\zeta$ is the factor converting the calcium current to calcium concentration change per unit time and $\beta_{\text{NMDAR}}, \beta_{\text{CaV}}$ is the fast buffering factor of calcium influx through NMDAR and CaV channels, respectively. We do not model the extrusion, diffusion, and slow buffering of calcium in detail, but describe them by "leak" term with single exponential decay to the resting calcium concentration $[Ca^{2+}]_0$ with the time constant $\tau_{Ca}$ equal to 12 ms.

***Calcium influx.***

**Calcium through NMDAR** The NMDA receptor-mediated calcium current is given by

$$I_{\text{NMDAR}} = g_{\text{NMDAR}} s_{\text{NMDAR}} B(V)\left(V - E_{\text{NMDAR,Ca}}\right). \quad [39]$$

$s_{\text{NMDAR}}$ shares the same value as in Eq. (28) while the reversal potential for calcium current $E_{\text{NMDAR,Ca}}$ is 140 mV instead of 0 mV for $E_{\text{NMDAR}}$ in Eq. (28).

**Calcium through voltage-gated calcium channels (CaV)** The calcium current through CaV channels is fully described above since the voltage-gated calcium channel is only permeable to calcium ions.

***Calcium trace with noise.*** The stochastic simulations are achieved by adding noise to the calcium dynamic during the synaptic events, through the stochastic opening and closing of ion channels. To represent this stochastic effect, we include two sources of noise following (2): (i) random maximum conductance of NMDAR at each presynaptic spike and (ii) random maximum conductance of the CaV channels at each postsynaptic spike and plateau potential. During each synaptic event, the channel conductance is drawn from a Gaussian distribution with a given mean and variance, chosen so that the stochastic system can reproduce smooth results, consistent with experiments. The same magnitude of noise as (2) is added for the simulation of "priming" and BTSP scenario, while a smaller magnitude of noise is used for STDP simulation (as shown in Table.S1) to get rid of a second "LTD" window for large negative $\Delta t$.

**Biochemical Reactions in a Single Chamber.** LTP/LTD follows from biochemical reactions in the PSD, which we model as a single chamber. These biochemical reactions describe the calcium-induced activation of both the kinase (CaMKII) and the phosphatase and the mutual inhibition between the two. The detailed reaction scheme is illustrated in Fig.S4. The whole system is initiated with calcium influx through NMDAR and CaV channels, activating CaMKII and phosphatase. The activated CaMKII and phosphatase will promote their own activation (self-activation) and inhibit each other through phosphorylation and dephosphorylation, following the idea of (9). Our detailed model of the phosphorylation of CaMKII is based on (2). Our model of the dephosphorylation of the phosphatase is more phenomological, similar to that in (9).

***Calcium binding with Calmodulin.*** Calmodulin is a calcium-binding messenger protein with two lobes (the N- and C- domains), and each lobe can bind to at most two calcium ions. The resulting calcium-calmodulin complex can further bind to CaMKII and trigger its activation. The binding of calcium ions acts cooperatively on each lobe. For simplicity, we only consider the number of calcium ions bound to calmodulin, without distinguishing between states with different binding sites but the same number of calcium ions bound. The reaction scheme is given as

$$\mathrm{C_0} \underset{-\mathrm{Ca^{2+}}}{\overset{\mathrm{K_1},\ +\mathrm{Ca^{2+}}}{\rightleftharpoons}} \mathrm{C_1} \underset{-\mathrm{Ca^{2+}}}{\overset{\mathrm{K_2},\ +\mathrm{Ca^{2+}}}{\rightleftharpoons}} \mathrm{C_2} \underset{-\mathrm{Ca^{2+}}}{\overset{\mathrm{K_3},\ +\mathrm{Ca^{2+}}}{\rightleftharpoons}} \mathrm{C_3} \underset{-\mathrm{Ca^{2+}}}{\overset{\mathrm{K_4},\ +\mathrm{Ca^{2+}}}{\rightleftharpoons}} \mathrm{C_4} \quad [40]$$

where $C_i$ is the Calmodulin with i calcium ions bounded. Since the binding and disassociation of calcium by calmodulin is much faster than the phosphorylation and dephosphorylation (10, 11), we introduce a 'fast-equilibrium' assumption where the calmodulin bound with different numbers of calcium ions are in equilibrium with each other and with the free calcium ions at each time point. This leads to the following four(4) algebraic equations ($[x]$ denotes the concentration of $x$):

$$K_1[C_1] = [Ca^{2+}][C_0],\ K_2[C_2] = [Ca^{2+}][C_1],\ K_3[C_3] = [Ca^{2+}][C_2],\ K_4[C_4] = [Ca^{2+}][C_3]. \quad [41]$$

This system can be solved for $[C_i], i = 0, ..., 3$ as a function of $[C_4]$:

$$[C_0] = \frac{K_1K_2K_3K_4}{[Ca^{2+}]^4}[C_4],\ [C_1] = \frac{K_2K_3K_4}{[Ca^{2+}]^3}[C_4],\ [C_2] = \frac{K_3K_4}{[Ca^{2+}]^2}[C_4],\ [C_3] = \frac{K_4}{[Ca^{2+}]}[C_4]. \quad [42]$$

This in turn yields an expression between $[C_4]$ and the concentration of *total* available calmodulin:

$$\sum_{i=0}^{4}[C_i] = \left(1 + \frac{K_4}{[Ca^{2+}]} + \frac{K_3K_4}{[Ca^{2+}]^2} + \frac{K_2K_3K_4}{[Ca^{2+}]^3} + \frac{K_1K_2K_3K_4}{[Ca^{2+}]^4}\right)[C_4] := K\left([Ca^{2+}]\right)[C_4]. \quad [43]$$

where $K\left([Ca^{2+}]\right)$ depends only on $[Ca^{2+}]$. We assume that, for the sequential phosphorylation, only $C_4$ (calmodulin bound with four ions) can bind to CaMKII. (Although there is evidence that calmodulin bound with two or three calcium ions can bind to CaMKII and promote its phosphorylation, the reported reaction rates are much smaller compared with calmodulin bound with four calcium ions (12).) .

***Binding and dissociation of molecules with CaMKII.*** CaMKII is a dodecameric holoenzyme, which is composed of two stacked hexameric rings formed by six functionally coupled subunits (13). Each subunit provides multiple binding sites for different proteins, and the binding of proteins can stimulate the phosphorylation or dephosphorylation of the subunits, as illustrated in Fig.S4(b). We assume that the binding and dissociation of proteins with CaMKII is in equilibrium, similar to calcium binding to Calmodulin.

For the family of non-phosphorylated subunits (chemicals with $K \cdot *$), we have that

$$K_{K\cdot pP}[K \cdot pP] = [K] \cdot [pP],\ K_{K\cdot ATP}[K \cdot ATP] = [K] \cdot [ATP],\ K_{K\cdot C_4}[K \cdot C_4] = [K] \cdot [C_4], \tag{44}$$

$$K_{K\cdot ATP\cdot C_4}[K \cdot C_4 \cdot ATP] = [K \cdot ATP] \cdot [C_4],\ K_{K\cdot C_4\cdot ATP}[K \cdot C_4 \cdot ATP] = [K \cdot C_4] \cdot [ATP]. \tag{45}$$

where $K$ denotes the non-phosphorylated subunit with no other proteins bound to it, and $K \cdot *$ is the non-phosphorylated subunit bound with $*$ $(* = ATP, pP, C_4, C_4 \cdot ATP)$. $K_x$ is the corresponding equilibrium constant which satisfies detailed balance,

$$K_{K\cdot ATP}K_{K\cdot ATP\cdot C_4} = K_{K\cdot C_4}K_{K\cdot C_4\cdot ATP}, \tag{46}$$

and guarantees the two equations in Eq. (45) are equivalent.

The probability $p_{K\cdot *}$ that the non-phosphorylated subunits are in the corresponding state $(K \cdot *)$ is given by

$$p_{K\cdot *} = \frac{[K \cdot *]}{[K] + [K \cdot ATP] + [K \cdot pP] + [K \cdot ATP] + [K \cdot C_4] + [K \cdot C_4 \cdot ATP]}. \tag{47}$$

Taking the fast-equilibrium into account, these probabilities satisfy

$$p_K = \frac{1}{1 + \frac{[ATP]}{K_{K\cdot ATP}} + \frac{[pP]}{K_{K\cdot pP}} + \frac{[C_4]}{K_{K\cdot C_4}} + \frac{[ATP][C_4]}{K_{K\cdot C_4}K_{K\cdot C_4\cdot ATP}}}. \tag{48}$$

$$p_{K\cdot *} = \frac{\frac{[*]}{K_{K\cdot *}}}{1 + \frac{[ATP]}{K_{K\cdot ATP}} + \frac{[pP]}{K_{K\cdot pP}} + \frac{[C_4]}{K_{K\cdot C_4}} + \frac{[ATP][C_4]}{K_{K\cdot C_4}K_{K\cdot C_4\cdot ATP}}}, * = pP, ATP, C_4. \tag{49}$$

$$p_{K\cdot C_4\cdot ATP} = \frac{\frac{[ATP][C_4]}{K_{K\cdot C_4}K_{K\cdot C_4\cdot ATP}}}{1 + \frac{[ATP]}{K_{K\cdot ATP}} + \frac{[pP]}{K_{K\cdot pP}} + \frac{[C_4]}{K_{K\cdot C_4}} + \frac{[ATP][C_4]}{K_{K\cdot C_4}K_{K\cdot C_4\cdot ATP}}}, \tag{50}$$

which can be determined when the values of $[pP], [C_4]$ are known.

The family of phosphorylated subunits $(pK \cdot *)$ satisfy similar fast-equilibrium relationships:

$$K_{pK\cdot P}[pK \cdot P] = [pK] \cdot [P],\ K_{pK\cdot ADP}[pK \cdot ADP] = [pK] \cdot [ADP],\ K_{pK\cdot C_4}[pK \cdot C_4] = [pK] \cdot [C_4], \tag{51}$$

$$K_{pK\cdot ADP\cdot C_4}[pK \cdot C_4 \cdot ADP] = [pK \cdot ADP] \cdot [C_4],\ K_{pK\cdot C_4\cdot ADP}[pK \cdot C_4 \cdot ADP] = [pK \cdot C_4] \cdot [ADP]. \tag{52}$$

where $pK$ is the phosphorylated subunit with no other proteins bounded and $pK \cdot *$ is the phosphorylated subunit bound with $*$ $(* = ADP, P, C_4, C_4 \cdot ADP)$. The equilibrium constants satisfy

$$K_{pK\cdot ADP}K_{pK\cdot ADP\cdot C_4} = K_{pK\cdot C_4}K_{pK\cdot C_4\cdot ADP}, \tag{53}$$

which ensures the equivalence of the two equations in Eq. (52).

The probability $p_{pK\cdot *}$ that the phosphorylated subunits are in the corresponding state $(pK \cdot *)$ is given by

$$p_{pK\cdot *} = \frac{[pK \cdot *]}{[pK] + [pK \cdot ADP] + [pK \cdot P] + [pK \cdot C_4] + [pK \cdot C_4 \cdot ADP]}. \tag{54}$$

With fast equilibrium, we have

$$p_{pK} = \frac{1}{1 + \frac{[ADP]}{K_{pK\cdot ADP}} + \frac{[P]}{K_{pK\cdot P}} + \frac{[C_4]}{K_{pK\cdot C_4}} + \frac{[ADP][C_4]}{K_{pK\cdot C_4}K_{pK\cdot C_4\cdot ADP}}}. \tag{55}$$

$$p_{pK\cdot *} = \frac{\frac{[*]}{K_{pK\cdot *}}}{1 + \frac{[ADP]}{K_{pK\cdot ADP}} + \frac{[P]}{K_{pK\cdot P}} + \frac{[C_4]}{K_{pK\cdot C_4}} + \frac{[ADP][C_4]}{K_{pK\cdot C_4}K_{pK\cdot C_4\cdot ADP}}}, * = P, ADP, C_4. \tag{56}$$

$$p_{pK\cdot C_4\cdot ADP} = \frac{\frac{[ADP][C_4]}{K_{pK\cdot C_4}K_{pK\cdot C_4\cdot ADP}}}{1 + \frac{[ADP]}{K_{pK\cdot ADP}} + \frac{[P]}{K_{pK\cdot P}} + \frac{[C_4]}{K_{pK\cdot C_4}} + \frac{[ADP][C_4]}{K_{pK\cdot C_4}K_{pK\cdot C_4\cdot ADP}}}. \tag{57}$$

which can be determined when the values of $[P], [C_4]$ are known.

Conservation laws must be included to complete the equilibrium equations. The dynamics of calmodulin are governed by the conservation law:

$$[CaM]_0 = [C_0] + [C_1] + [C_2] + [C_3] + [C_4] + [K \cdot C_4] + [pK \cdot C_4] + [K \cdot C_4 \cdot ATP] + [pK \cdot C_4 \cdot ADP]. \tag{58}$$

where $[CaM]_0$ denotes the total concentration of calmodulin in the whole system. At each time point, the concentration of the phosphorylated and unphosphorylated subunits are determined ($[K]_0$ and $[pK]_0$). Combined with the conversation law of the phosphatase $[P]$, we have four more equations:

$$[K]_0 = [K] + [K \cdot pP] + [K \cdot ATP] + [K \cdot C_4] + [K \cdot C_4 \cdot ATP]. \quad [59]$$

$$[pK]_0 = [pK] + [pK \cdot P] + [pK \cdot ADP] + [pK \cdot C_4] + [pK \cdot C_4 \cdot ADP]. \quad [60]$$

$$[P]_0 = [pK \cdot P] + [P]. \quad [61]$$

$$[pP]_0 = [K \cdot pP] + [pP]. \quad [62]$$

Eq. (58) to Eq. (62) form a nonlinear system with five equations on five independent variables: $[C_4], [K], [pK], [P], [pP]$, such that it is solvable. The detailed method to solve the nonlinear system is presented in the 'Numerical Methods' section.

***Phosphorylation and Dephosphorylation of CaMKII.*** The binding of calcium-calmodulin complex will initiate the phosphorylation at threonine-286 (Thr-286) in the regulatory domain of CaMKII, which disrupts the binding of the regulatory domain to the kinase domain (13). After being phosphorylated, CaMKII subunits have a stronger affinity for calcium-calmodulin (14) and can stimulate the phosphorylation of their neighboring subunits, which prolongs the activation after the calcium stimulus (15). They can also bind to and phosphorylate other enzymatic substrates like AMPAR and NMDAR, leading to synapse potentiation (16, 17). CaMKII activity is treated in detail in the model, and it serves as an indicator of potentiation, with some aspects not considered for simplicity, such as different isoforms of CaMKII subunits or the phosphorylation at Thr-305 and 306.

In modelling the phosphorylation and dephosphorylation of CaMKII, we follow closely Graupner, and Brunel (2). Since little is known about how the two rings of the CaMKII interact and how that affects phosphorylation (18), we suppose the two are independent of each other and model only one six-subunit ring. As each subunit can be either phosphorylated or unphosphorylated, there are 14 phosphorylated states for a six-subunit ring under the rotation invariance. We denote them as open states $O^{(i)}, 1 \leq i \leq 14$ (ordered by the number of phosphorylated subunits), as shown in Fig.S4(d). The transition between these different states can be divided into two categories: (i) the spontaneous phosphorylation and dephosphorylation of one subunit and (ii) the neighboring phosphorylation of CaMKII (and its reversal reaction).

We assume that the *spontaneous* phosphorylation and dephosphorylation follow the following rules: (i) one unphosphorylated subunit can only become phosphorylated with rate $\alpha_K$ when bound to ATP or with rate $\alpha_{KC}$ when bound to both ATP and calmodulin. The rate for the corresponding reversal reaction is $\beta_K$ and $\beta_{KC}$. (ii) one phosphorylated subunit can only become dephosphorylated with rate $\alpha_P$ when bound to active phosphatase, and the corresponding reversal reaction has rate $\beta_P$. Taking the probability of protein bounding into account, we know the net rates for one subunit's spontaneous phosphorylation and dephosphorylation equal to

$$\alpha = \alpha_K \cdot p_{K \cdot ATP} + \alpha_{KC} \cdot p_{K \cdot C_4 \cdot ATP} + \beta_P \cdot p_{K \cdot pP}, \quad \beta = \beta_K \cdot p_{pK \cdot ADP} + \beta_{KC} \cdot p_{pk \cdot C_4 \cdot ADP} + \alpha_P \cdot p_{pK \cdot P} \cdot \left(1 + \frac{[P]_b}{[P]}\right), \quad [63]$$

where $[P]_b$ is the baseline activity of phosphatase similar to the formula in (9); while the matrices of transition between 14 states of CaMKII are given by

$$S_1 = \begin{pmatrix} -6 & 0 & 0 & 0 & 0 & 0 & 0 & 0 & 0 & 0 & 0 & 0 & 0 & 0 \\ 6 & -5 & 0 & 0 & 0 & 0 & 0 & 0 & 0 & 0 & 0 & 0 & 0 & 0 \\ 0 & 2 & -4 & 0 & 0 & 0 & 0 & 0 & 0 & 0 & 0 & 0 & 0 & 0 \\ 0 & 2 & 0 & -4 & 0 & 0 & 0 & 0 & 0 & 0 & 0 & 0 & 0 & 0 \\ 0 & 1 & 0 & 0 & -4 & 0 & 0 & 0 & 0 & 0 & 0 & 0 & 0 & 0 \\ 0 & 0 & 2 & 1 & 0 & -3 & 0 & 0 & 0 & 0 & 0 & 0 & 0 & 0 \\ 0 & 0 & 1 & 1 & 2 & 0 & -3 & 0 & 0 & 0 & 0 & 0 & 0 & 0 \\ 0 & 0 & 1 & 1 & 2 & 0 & 0 & -3 & 0 & 0 & 0 & 0 & 0 & 0 \\ 0 & 0 & 0 & 1 & 0 & 0 & 0 & 0 & -3 & 0 & 0 & 0 & 0 & 0 \\ 0 & 0 & 0 & 0 & 0 & 2 & 1 & 1 & 0 & -2 & 0 & 0 & 0 & 0 \\ 0 & 0 & 0 & 0 & 0 & 1 & 1 & 1 & 3 & 0 & -2 & 0 & 0 & 0 \\ 0 & 0 & 0 & 0 & 0 & 0 & 1 & 1 & 0 & 0 & 0 & -2 & 0 & 0 \\ 0 & 0 & 0 & 0 & 0 & 0 & 0 & 0 & 0 & 2 & 2 & 2 & -1 & 0 \\ 0 & 0 & 0 & 0 & 0 & 0 & 0 & 0 & 0 & 0 & 0 & 0 & 1 & 0 \end{pmatrix} \text{(spontaneous phosphorylation)}, \qquad [64]$$

$$S_2 = \begin{pmatrix} 0 & 1 & 0 & 0 & 0 & 0 & 0 & 0 & 0 & 0 & 0 & 0 & 0 & 0 \\ 0 & -1 & 2 & 2 & 2 & 0 & 0 & 0 & 0 & 0 & 0 & 0 & 0 & 0 \\ 0 & 0 & -2 & 0 & 0 & 2 & 1 & 1 & 0 & 0 & 0 & 0 & 0 & 0 \\ 0 & 0 & 0 & -2 & 0 & 1 & 1 & 1 & 3 & 0 & 0 & 0 & 0 & 0 \\ 0 & 0 & 0 & 0 & -2 & 0 & 1 & 1 & 0 & 0 & 0 & 0 & 0 & 0 \\ 0 & 0 & 0 & 0 & 0 & -3 & 0 & 0 & 0 & 2 & 1 & 0 & 0 & 0 \\ 0 & 0 & 0 & 0 & 0 & 0 & -3 & 0 & 0 & 1 & 1 & 2 & 0 & 0 \\ 0 & 0 & 0 & 0 & 0 & 0 & 0 & -3 & 0 & 1 & 1 & 2 & 0 & 0 \\ 0 & 0 & 0 & 0 & 0 & 0 & 0 & 0 & -3 & 0 & 1 & 0 & 0 & 0 \\ 0 & 0 & 0 & 0 & 0 & 0 & 0 & 0 & 0 & -4 & 0 & 0 & 2 & 0 \\ 0 & 0 & 0 & 0 & 0 & 0 & 0 & 0 & 0 & 0 & -4 & 0 & 2 & 0 \\ 0 & 0 & 0 & 0 & 0 & 0 & 0 & 0 & 0 & 0 & 0 & -4 & 1 & 0 \\ 0 & 0 & 0 & 0 & 0 & 0 & 0 & 0 & 0 & 0 & 0 & 0 & -5 & 6 \\ 0 & 0 & 0 & 0 & 0 & 0 & 0 & 0 & 0 & 0 & 0 & 0 & 0 & -6 \end{pmatrix} \text{(spontaneous dephosphorylation)}, \qquad [65]$$

such that the net rate matrix is

$$S_A = \alpha S_1 + \beta S_2. \qquad [66]$$

The auto-phosphorylation mechanism plays an essential role in maintaining the activity of CaMKII after the disassociation of calcium-calmodulin. Auto-phosphorylation, also termed neighboring phosphorylation, describes the phenomenon of an intersubunit process during which one subunit acts as a substrate and the adjacent subunit as a catalyst. We only consider the case where the process proceeds in the counter-clockwise direction, as is common in the literature (2, 19). (There are no experimental observations to the contrary.) Binding with calmodulin is required for the substrate subunit to become phosphorylated (17). However, the neighboring substrate can catalyze the reaction wherever its auto-inhibitory domain is released by binding with calcium-calmodulin or phosphorylation at Thr-286 (13).

In *auto-phosphorylation*, we suppose that the neighboring phosphorylation satisfies the following rules: one unphosphorylated subunit bound to both ATP and calmodulin can become phosphorylated (i) with rate $\alpha_{K,NC}$ when its neighbor is unphosphorylated and bound to calmodulin, with reversal reaction rate $\beta_{K,NC}$; (ii) with rate $\alpha_{K,NP}$ when its neighbor is phosphorylated and not bound to phosphatase, and the corresponding reversal reaction has rate $\beta_{K,NP}$. Assuming the protein binding is independent of the neighboring subunits and taking the probability of fast-equilibrium and the transition between 14 states of CaMKII into account, net rates for neighboring phosphorylation are given by

$$S_{NP} = \alpha_{K,NC} \cdot (p_{K \cdot C_4} + p_{K \cdot C_4 \cdot ATP}) \, p_{K \cdot C_4 \cdot ATP} S_3 + \alpha_{K,NP} \cdot (1 - p_{pK \cdot P}) \, p_{K \cdot C_4 \cdot ATP} S_4, \qquad [67]$$

$$S_3 = \begin{pmatrix} -6 & 0 & 0 & 0 & 0 & 0 & 0 & 0 & 0 & 0 & 0 & 0 & 0 & 0 \\ 6 & -4 & 0 & 0 & 0 & 0 & 0 & 0 & 0 & 0 & 0 & 0 & 0 & 0 \\ 0 & 1 & -3 & 0 & 0 & 0 & 0 & 0 & 0 & 0 & 0 & 0 & 0 & 0 \\ 0 & 2 & 0 & -2 & 0 & 0 & 0 & 0 & 0 & 0 & 0 & 0 & 0 & 0 \\ 0 & 1 & 0 & 0 & -2 & 0 & 0 & 0 & 0 & 0 & 0 & 0 & 0 & 0 \\ 0 & 0 & 1 & 0 & 0 & -2 & 0 & 0 & 0 & 0 & 0 & 0 & 0 & 0 \\ 0 & 0 & 1 & 1 & 0 & 0 & -1 & 0 & 0 & 0 & 0 & 0 & 0 & 0 \\ 0 & 0 & 1 & 0 & 2 & 0 & 0 & -1 & 0 & 0 & 0 & 0 & 0 & 0 \\ 0 & 0 & 0 & 1 & 0 & 0 & 0 & 0 & 0 & 0 & 0 & 0 & 0 & 0 \\ 0 & 0 & 0 & 0 & 0 & 1 & 0 & 0 & 0 & -1 & 0 & 0 & 0 & 0 \\ 0 & 0 & 0 & 0 & 0 & 1 & 1 & 0 & 0 & 0 & 0 & 0 & 0 & 0 \\ 0 & 0 & 0 & 0 & 0 & 0 & 0 & 1 & 0 & 0 & 0 & 0 & 0 & 0 \\ 0 & 0 & 0 & 0 & 0 & 0 & 0 & 0 & 0 & 1 & 0 & 0 & 0 & 0 \\ 0 & 0 & 0 & 0 & 0 & 0 & 0 & 0 & 0 & 0 & 0 & 0 & 0 & 0 \end{pmatrix} \text{(neighbor with calmodulin)}, \quad [68]$$

$$S_4 = \begin{pmatrix} 0 & 0 & 0 & 0 & 0 & 0 & 0 & 0 & 0 & 0 & 0 & 0 & 0 & 0 \\ 0 & -1 & 0 & 0 & 0 & 0 & 0 & 0 & 0 & 0 & 0 & 0 & 0 & 0 \\ 0 & 1 & -1 & 0 & 0 & 0 & 0 & 0 & 0 & 0 & 0 & 0 & 0 & 0 \\ 0 & 0 & 0 & -2 & 0 & 0 & 0 & 0 & 0 & 0 & 0 & 0 & 0 & 0 \\ 0 & 0 & 0 & 0 & -2 & 0 & 0 & 0 & 0 & 0 & 0 & 0 & 0 & 0 \\ 0 & 0 & 1 & 1 & 0 & -1 & 0 & 0 & 0 & 0 & 0 & 0 & 0 & 0 \\ 0 & 0 & 0 & 0 & 2 & 0 & -2 & 0 & 0 & 0 & 0 & 0 & 0 & 0 \\ 0 & 0 & 0 & 1 & 0 & 0 & 0 & -2 & 0 & 0 & 0 & 0 & 0 & 0 \\ 0 & 0 & 0 & 0 & 0 & 0 & 0 & 0 & -3 & 0 & 0 & 0 & 0 & 0 \\ 0 & 0 & 0 & 0 & 0 & 1 & 1 & 1 & 0 & -1 & 0 & 0 & 0 & 0 \\ 0 & 0 & 0 & 0 & 0 & 0 & 0 & 1 & 3 & 0 & -2 & 0 & 0 & 0 \\ 0 & 0 & 0 & 0 & 0 & 0 & 1 & 0 & 0 & 0 & 0 & -2 & 0 & 0 \\ 0 & 0 & 0 & 0 & 0 & 0 & 0 & 0 & 0 & 1 & 2 & 2 & -1 & 0 \\ 0 & 0 & 0 & 0 & 0 & 0 & 0 & 0 & 0 & 0 & 0 & 0 & 1 & 0 \end{pmatrix} \text{(neighbor phosphorylated)}, \quad [69]$$

and the net rates for the reversal reactions is given by

$$S_{NR} = \beta_{K,NC} \cdot (p_{K \cdot C_4} + p_{K \cdot C_4 \cdot ATP})\, p_{pK \cdot C_4 \cdot ADP} S_5 + \beta_{K,NP} \cdot (1 - p_{pK \cdot P})\, p_{K \cdot C_4 \cdot ATP} S_6, \quad [70]$$

$$S_5 = \begin{pmatrix} 0 & 1 & 0 & 0 & 0 & 0 & 0 & 0 & 0 & 0 & 0 & 0 & 0 & 0 \\ 0 & -1 & 1 & 2 & 2 & 0 & 0 & 0 & 0 & 0 & 0 & 0 & 0 & 0 \\ 0 & 0 & -1 & 0 & 0 & 1 & 1 & 1 & 0 & 0 & 0 & 0 & 0 & 0 \\ 0 & 0 & 0 & -2 & 0 & 0 & 1 & 0 & 3 & 0 & 0 & 0 & 0 & 0 \\ 0 & 0 & 0 & 0 & -2 & 0 & 0 & 1 & 0 & 0 & 0 & 0 & 0 & 0 \\ 0 & 0 & 0 & 0 & 0 & -1 & 0 & 0 & 0 & 1 & 1 & 0 & 0 & 0 \\ 0 & 0 & 0 & 0 & 0 & 0 & -2 & 0 & 0 & 0 & 1 & 0 & 0 & 0 \\ 0 & 0 & 0 & 0 & 0 & 0 & 0 & -2 & 0 & 0 & 0 & 2 & 0 & 0 \\ 0 & 0 & 0 & 0 & 0 & 0 & 0 & 0 & -3 & 0 & 0 & 0 & 0 & 0 \\ 0 & 0 & 0 & 0 & 0 & 0 & 0 & 0 & 0 & -1 & 0 & 0 & 1 & 0 \\ 0 & 0 & 0 & 0 & 0 & 0 & 0 & 0 & 0 & 0 & -2 & 0 & 0 & 0 \\ 0 & 0 & 0 & 0 & 0 & 0 & 0 & 0 & 0 & 0 & 0 & -2 & 0 & 0 \\ 0 & 0 & 0 & 0 & 0 & 0 & 0 & 0 & 0 & 0 & 0 & 0 & -1 & 0 \\ 0 & 0 & 0 & 0 & 0 & 0 & 0 & 0 & 0 & 0 & 0 & 0 & 0 & 0 \end{pmatrix} \text{(neighbor with calmodulin)}, \quad [71]$$

$$S_6 = \begin{pmatrix} 0 & 0 & 0 & 0 & 0 & 0 & 0 & 0 & 0 & 0 & 0 & 0 & 0 & 0 \\ 0 & 0 & 1 & 0 & 0 & 0 & 0 & 0 & 0 & 0 & 0 & 0 & 0 & 0 \\ 0 & 0 & -1 & 0 & 0 & 1 & 0 & 0 & 0 & 0 & 0 & 0 & 0 & 0 \\ 0 & 0 & 0 & 0 & 0 & 1 & 0 & 1 & 0 & 0 & 0 & 0 & 0 & 0 \\ 0 & 0 & 0 & 0 & 0 & 0 & 1 & 0 & 0 & 0 & 0 & 0 & 0 & 0 \\ 0 & 0 & 0 & 0 & 0 & -2 & 0 & 0 & 0 & 1 & 0 & 0 & 0 & 0 \\ 0 & 0 & 0 & 0 & 0 & 0 & -1 & 0 & 0 & 1 & 0 & 2 & 0 & 0 \\ 0 & 0 & 0 & 0 & 0 & 0 & 0 & -1 & 0 & 1 & 1 & 0 & 0 & 0 \\ 0 & 0 & 0 & 0 & 0 & 0 & 0 & 0 & 0 & 0 & 1 & 0 & 0 & 0 \\ 0 & 0 & 0 & 0 & 0 & 0 & 0 & 0 & 0 & -3 & 0 & 0 & 1 & 0 \\ 0 & 0 & 0 & 0 & 0 & 0 & 0 & 0 & 0 & 0 & -2 & 0 & 2 & 0 \\ 0 & 0 & 0 & 0 & 0 & 0 & 0 & 0 & 0 & 0 & 0 & -2 & 1 & 0 \\ 0 & 0 & 0 & 0 & 0 & 0 & 0 & 0 & 0 & 0 & 0 & 0 & -4 & 6 \\ 0 & 0 & 0 & 0 & 0 & 0 & 0 & 0 & 0 & 0 & 0 & 0 & 0 & -6 \end{pmatrix} \text{(neighbor phosphorylated)}, \quad [72]$$

Combining the equations above, the dynamical system governing phosphorylation satisfies the following equation:

$$\frac{d[O^{(i)}]}{dt} = S(i,:)\ \mathbf{O}, \quad 1 \le i \le 14, \quad S = S_A + S_{NP} + S_{NR}. \qquad [73]$$

***Docking and undocking of CaMKII.*** The CaMKII holoenzyme exists in both compact and extended auto-inhibited states (20, 21). The compact state can be bound to actin filaments in the spine head and dendritic shaft, with the compact state's release from the actin is triggered by calcium-calmodulin binding (22). In our model, we include this compact state 'C' as a $15^{th}$ state of CaMKII. CaMKII in the compact state can become open with a calcium-dependent rate, and the CaMKII in the extended state can return to the compact state when it has no phosphorylated subunits and is not bound to calmodulin or phosphatase. The transition between the compact state and extended state is given by

$$\frac{d[C]}{dt} = -B\left(\left[Ca^{2+}\right]\right)\mu[C] + \left(p_K + p_{K\cdot ATP}\right)^n \nu[O^{(1)}], \qquad [74]$$

$$B\left([Ca^{2+}]\right) = \frac{\left[Ca^{2+}\right]^2}{[Ca^{2+}]^2 + 0.2^2}, \qquad [75]$$

where $[C]$ and $[O^{(1)}]$ are the concentration of the CaMKII in the compact state and the extended state with no phosphorylated subunits, respectively, and $p_K + p_{K\cdot ATP}$ is the probability that one unphosphorylated subunit is not bound to calmodulin or phosphatase.

***Full CaMKII Dynamic.*** With the above notation of the matrix $S$ and variables of the open states $\mathbf{O} = \left([O^{(1)}], \cdots, [O^{(14)}]\right)^t$, the dynamics of the system is given by

$$\begin{aligned}
\frac{d[C]}{dt} &= -B\left([Ca^{2+}]\right)\mu[C] + \left(p_K + p_{K\cdot ATP}\right)^n \nu[O^{(1)}], \\
\frac{d[O^{(1)}]}{dt} &= B\left([Ca^{2+}]\right)\mu[C] - \left(p_K + p_{K\cdot ATP}\right)^n \nu[O^{(1)}] + S(1,:)\ \mathbf{O}, \\
\frac{d[O^{(i)}]}{dt} &= S(i,:)\ \mathbf{O}, \quad 1 < i \le 14. \qquad [76]
\end{aligned}$$

We have $[K]_0 = \sum_i \mathbf{K}(i)[O^{(i)}] = \mathbf{K}\cdot\mathbf{O}$, $[pK]_0 = \sum_i \mathbf{pK}(i)[O^{(i)}] = \mathbf{pK}\cdot\mathbf{O}$ and the conservation law for CaMKII: $[C] + ([K]_0 + [pK]_0)/n = [\text{CaMKII}]_0$. Details about the method to simulate the system is discussed in the 'Numerical Methods' section.

***Phenomenological phosphatase dynamics.*** Little seems to be known experimentally about how phosphatase is activated by calcium influx (23) Here we adapt a phenomenological model from (9) of the dynamics of the phosphatase concentration:

$$\frac{d[P]_0}{dt} = \alpha_P^{(\text{auto})}\frac{[pP]_0}{K_{\text{m50,pP}} + [pP]_0}[P] - \alpha_P[pK\cdot P] + \beta_P[K\cdot pP] + r_{\text{base}} + \alpha_{PC}Q\left([Ca^{2+}]\right)[pP]_0. \qquad [77]$$

with $[P]_0 + [pP]_0 = [P]_{tot},\ [P] + [pK\cdot P] = [P]_0,\ [pK\cdot P] = p_{pK\cdot P}[pK]_0$ and

$$Q\left([Ca^{2+}]\right) = \frac{\left[[Ca^{2+}] - 0.1\right]_+^2}{[[Ca^{2+}] - 0.1]_+^2 + 0.09^2} + 0.005, \quad [x]_+ = \max(x, 0). \qquad [78]$$

$[P]_0$ and $[pP]_0$ are the concentrations of active (dephosphorylated) and inactive (phosphorylated) phosphatase, respectively and $r_{\text{base}}$ is the baseline dephosphorylation activity rate. $\alpha_{PC}Q\left([Ca^{2+}]\right)$ describes the calcium dependent rate of dephosphorylation with a lower activation threshold (compared with CaMKII).

***AMPAR trafficking.*** An idealized model of AMPAR trafficking is taken from (9) to provide a readout mechanism for the CaMKII and phosphatase system. AMPAR is inserted into and removed from the membrane with the corresponding rates $k_1$ and $k_2$, described with first-order dynamics:

$$\frac{d}{dt}A = k_1\left(A_{tot} - A\right) - k_2 A, \qquad [79]$$

where $A$ is the number of AMPA receptors on the membrane, and $A_{tot}$ is the number of total available AMPA receptors, which is a constant number during the simulation. The insertion rate and removal rate depend on the activity of CaMKII and phosphatase, respectively. The dependency takes simply the formula of a linear function:

$$k_1 = c_1[pK]_0 + c_3, \qquad [80]$$

$$k_2 = c_2[P]_0 + c_4, \qquad [81]$$

where $c_1, c_2$ are the scaling factors, and $c_3$ and $c_4$ are the insertion and removal rates independent of CaMKII and phosphatase activities, respectively. We suppose that the AMPAR trafficking operates in a much slower timescale such that there will be no feedback effect of the trafficking on the magnitude of AMPAR current during the stimulus protocol.

**Model Parameters.** The parameters for the leaky integrate-and-fire neuron and the calcium signaling in the spine head are given in Table.S1 and the parameters for the biochemical reactions involving CaMKII & PP are given in Table.S2. Most of the parameters are taken from previous literature, while some of them are modified/refitted to obtain results consistent with previous experiments/modeling works.

***Parameters of phenomenological phosphatase dynamics.*** Following the general concept of coupled phosphatase and kinase switch from Pi & Lisman (9), the parameters for the phenomenological PP pathway are chosen to be compatible with the CaMKII pathway, leading to a realistic tri-stable system. A refitting of parameters of the PP pathway is required since our CaMKII pathway is modeled with a more biologically realistic description instead of the phenomenological model in (9).

The concentration of PP is chosen to be comparable with CaMKII subunits, same relationship as in (9). Given that CaMKII has a total concentration of $18\mu M$ and each CaMKII consists of six subunits (in our model), we choose the concentration of PP to be $100\mu M$. The phenomenological PP in our model could be a mixed group of subtypes of phosphatase (PP1, PP2A, PP2B); thus, the concentration is probably a summation of all these subtypes. Moreover, the reaction rates are chosen, satisfying two criteria: i) producing a tri-stable system and ii) leading to a realistic response. We fine-tune the parameters such that the system can maintain five steady states with three of them stable (the nullclines are shown in Fig.S5A) based on the phase plane analysis. However, the nullclines can only constrain the relative magnitude of the rates (as speeding up/down the rates together will not affect the shape of the nullcline). We further determine the velocity of the phosphatase dynamics based on how it matches with CaMKII dynamics and reproduces realistic responses (especially for STDP). Parameters tuned following the above procedure are given in Table.S2.

### Numerical Methods.

***Nonlinear Equation Solver.*** A system of coupled nonlinear equations emerging from the quasi-equilibrium assumptions must be solved at each step of the numerical integration. We have reduced the system of nonlinear equations to a single, uni-variate equation to alleviate the complexity of solving nonlinear systems.

The original equations can be simplified as

$$[K]_0 = [K]\left[1+\frac{[ATP]}{K_{K\cdot ATP}}+\frac{[pP]_0}{K_{K\cdot pP}+[K]}+\frac{[C_4]}{K_{K\cdot C_4}}\left(1+\frac{[ATP]}{K_{K\cdot C_4\cdot ATP}}\right)\right], \quad [82]$$

$$[pK]_0 = [pK]\left[1+\frac{[ADP]}{K_{pK\cdot ADP}}+\frac{[P]_0}{K_{pK\cdot P}+[pK]}+\frac{[C_4]}{K_{pK\cdot C_4}}\left(1+\frac{[ADP]}{K_{pK\cdot C_4\cdot ADP}}\right)\right], \quad [83]$$

$$[C_4] = \frac{[CaM]_0}{K([Ca^{2+}])+\left(1+\frac{[ATP]}{K_{K\cdot C_4\cdot ATP}}\right)\frac{[K]}{K_{K\cdot C_4}}+\left(1+\frac{[ADP]}{K_{pK\cdot C_4\cdot ADP}}\right)\frac{[pK]}{K_{pK\cdot C_4}}}. \quad [84]$$

Eq. (82) and Eq. (83) indicate that $[K]$ and $[pK]$ are both implicit functions of $[C_4]$ and take unique value in the biologically reasonable range. The system can then be reduced to a single nonlinear equation of $[C_4]$ after the relationship of $[K]$ and $[pK]$ with $[C_4]$ are plugged into Eq. (84). The value of $[C_4]$ is obtained by solving the single equation with a MATLAB built-in function 'fzero'. The value of $[K]$ and $[pK]$ can then be solved from Eq. (82) and Eq. (83) given the known value of $[C_4]$ and that fully solves the system.

***ODE Solver.*** The numerical simulation of the ordinary differential equations is implemented in MATLAB. Since the system involves chemical reactions with rates covering different scales, a stiff method with adaptive step size will be more stable and efficient. We use 'ode15s', a MATLAB built-in, variable order method for solving stiff differential equations to integrate the system numerically. We set the tolerance of relative error to be 1e-7 and the tolerance of absolute error to be 1e-9 for numerical stability.

We have introduced a piece-wise integration scheme, as the system is driven by the 'spike-nature' of the voltage and calcium dynamics. The whole time interval is divided into two categories of sub-intervals: wide sub-intervals with no synaptic events and narrow ones containing synaptic events. Max step size for numerical integration is set to be 100ms for wide sub-intervals and 1ms for narrow ones. That both allows efficiency and ensures any spike events will not be neglected. It takes approximately 2 minutes to run a 180-second simulation containing 60 presynaptic and 60 postsynaptic spikes on one AMD R9-5900HX laptop CPU thread.

**Table S1. Table of all parameters**

| Parameter | Definition | Value | Reference and/or Comment |
|---|---|---|---|
| $C_m$ | Whole membrane capacitance | $100\ pF$ | (2) |
| $g_{\text{Leak}}$ | Maximum leak conductance | $5\ nS$ | (2) |
| $E_{\text{Leak}}$ | Leaking potential | $-70\ mV$ | (24) |
| $g_{\text{Na}}$ | Maximum sodium conductance | $0.7\ \mu S$ | (2) |
| $E_{\text{Na}}$ | Sodium reversal potential | $60\ mV$ | (2) |
| $g_{\text{K}}$ | Maximum potassium conductance | $1.3\ \mu S$ | (2) |
| $E_{\text{K}}$ | Potassium reversal potential | $-80\ mV$ | (2) |
| $g_{\text{AMPAR}}$ | Maximal conductance of total AMPARs (per spine) | $19.5\ nS$ | (2) |
| $E_{\text{AMPAR}}$ | AMPA-mediated current reversal potential | $0\ mV$ | (2) |
| $\tau_{\text{AMPAR}}$ | Time constant for AMPA gating variable | $2\ ms$ | (2) |
| $\tau'_{\text{AMPAR}}$ | Time constant for presynaptic variable | $0.05\ ms$ | (2) |
| $\alpha_s$ | Effect magnitude of AMPA gating variable | $1\ /ms$ | (2) |
| $\alpha_x$ | Effect magnitude of presynaptic variable | $1$ | (2) |
| $N_{\text{NMDAR}}$ | Number of NMDAR (per spine) | $7$ | $5-30$ (25, 26) |
| $\bar{g}_{\text{NMDAR}}$ | Maximal conductance of single NMDAR | $53\ pS$ | $53 \pm 5\ pS$ (27) |
| $g_{\text{NMDAR}}$ | Maximal conductance of total NMDARs (per spine) | $0.371\ nS$ | $g_{\text{NMDAR}} = N_{\text{NMDAR}} \cdot \bar{g}_{\text{NMDAR}}$. |
| $\sigma_{\text{NMDAR}}$ | SD of the Gaussian noise added ("Priming", BTSP) | $3.3\%$ of $g_{\text{NMDAR}}$ | (2) |
| $\sigma_{\text{NMDAR}}$ | SD of the Gaussian noise added (STDP) | $1\%$ of $g_{\text{NMDAR}}$ | Fitting data from (28) |
| $E_{\text{NMDAR}}$ | NMDAR-mediated current reversal potential | $0\ mV$ | (2) |
| $E_{\text{NMDAR},Ca}$ | NMDAR-mediated Calcium reversal potential | $140\ mV$ | (2) |
| $[Mg^{2+}]$ | Extracellular Magnesium concentration | $1\ mM$ | (2) |
| $\tau_{\text{NMDAR}}$ | Time constant for NMDAR gating variable | $80\ ms$ | (2) |
| $\tau'_{\text{NMDAR}}$ | Time constant for presynaptic variable | $2\ ms$ | (2) |
| $\alpha_s$ | Effect magnitude of NMDAR gating variable | $1\ /ms$ | (2) |
| $\alpha_x$ | Effect magnitude of presynaptic variable | $1$ | (2) |
| $g_{\text{CaV1.2}}$ | Strength of Ca current flux due to CaV1.2 | $1.656\ mol/(cm\ C)$ | Fitting (2) |
| $g_{\text{CaV1.3}}$ | Strength of Ca current flux due to CaV1.3 | $0.404\ mol/(cm\ C)$ | Fitting (2) & the CaV1.2/CaV1.3 ratio (4:1) (29, 30) |
| $\tilde{g}_{\text{CaV1.2}}$ | Strength of local Ca current flux due to CaV1.2 | $82.82\ mol/(cm\ C)$ | $\tilde{g}_{\text{CaV1.2}} = 50 \cdot g_{\text{CaV1.2}}$ (based on (5)) |
| $\tilde{g}_{\text{CaV1.3}}$ | Strength of local Ca current flux due to CaV1.3 | $20.2\ mol/(cm\ C)$ | $\tilde{g}_{\text{CaV1.3}} = 50 \cdot g_{\text{CaV1.3}}$ (based on (5)) |
| $\sigma_{\text{CaV}}$ | SD of the Gaussian noise added ("Priming", BTSP) | $10\%$ of $g_{\text{CaV}}$ | (2) |
| $\sigma_{\text{CaV}}$ | SD of the Gaussian noise added ("Priming", BTSP) | $2\%$ of $g_{\text{CaV}}$ | Fitting data from (28) |
| $P_{\text{Ca}}$ | Constant | $0.00054\ cm/s$ | (5) |
| $F$ | Faraday constant | $96.5\ C/mmol$ | |
| $R$ | Universal gas constant | $8.315\ J/(mol * K)$ | |
| $T$ | Temperature | $308K$ | |
| $[Ca^{2+}]_o$ | External calcium concentration | $1800\ \mu M$ | (5) |
| $\tilde{c}_p$ | Threshold for Ca dependence | $3.0\ \mu M$ | (5) |
| $\tau_{po}$ | Time constant of activation | $1\ ms$ | (5) |
| $r_1$ | Opening rate | $0.3\ ms^{-1}$ | (5) |
| $r_2$ | Closing rate | $3\ ms^{-1}$ | (5) |
| $s'_1$ | Inactivation rate | $0.00195\ ms^{-1}$ | (5) |
| $k'_1$ | Inactivation rate | $0.00413\ ms^{-1}$ | (5) |
| $k_2$ | Inactivation rate | $0.0001\ ms^{-1}$ | (5) |
| $k'_{21}$ | Inactivation rate | $0.00224\ ms^{-1}$ | (5) |
| $T_{Ba}$ | Time constant | $450\ ms$ | (5) |
| $\Delta V$ (CaV1.2) | Displacement of voltage from CaV1.2 model | $0\ mV$ | Fitting data from (7, 8) |
| $\Delta V$ (CaV1.3) | Displacement of voltage from CaV1.2 model | $30\ mV$ | Fitting data from (7, 8) |
| $\tau_{Ca}$ | Time constant of calcium dynamics | $12\ ms$ | (2) |
| $[Ca^{2+}]_0$ | Resting calcium concentration in spine head | $0.094\ \mu M$ | $0.07 \pm 0.029\ \mu M$ (31) |
| $V_{\text{spine}}$ | Volume of spine head | $1\ \mu m^3$ | (2) |
| $\zeta$ | Factor converting calcium current to concentration | $5.18 \cdot 10^3\ \mu M/pC$ | $\zeta = 1/(2FV_{\text{spine}})$ |
| $\beta_{\text{NMDAR}}$ | Buffering factor of calcium through NMDAR | $1/1000$ | (2) |
| $\beta_{\text{CaV}}$ | Buffering factor of calcium through CaV | $1/100$ | (2) |

**Table S2. CaMKII & PP parameters**

| Parameter | Definition | Value | Reference and/or Comment |
|---|---|---|---|
| $[CaM]_0$ | Total concentration of Calmodulin | $30\mu M$ | (32) |
| $[CaMKII]_0$ | Total concentration of CaMKII | $18\mu M$ | Modified from $16.67\ \mu M$ in (2) |
| $[P]_{tot}$ | Total concentration of Phosphatase | $100\mu M$ | See text |
| $[ATP]$ | Available concentration of ATP | $1000\mu M$ | (33) |
| $[ADP]$ | Available concentration of ADP | $0.1\ \mu M$ | Chosen $\ll [ATP]$ for effective phosphorylation |
| $K_1$ | Dissociation constant of Calcium to CaM | $0.1\ \mu M$ | (2) |
| $K_2$ | Dissociation constant of Calcium to CaM | $0.02\ \mu M$ | Modified from $0.025\ \mu M$ in (2) |
| $K_3$ | Dissociation constant of Calcium to CaM | $5\ \mu M$ | (10) |
| $K_4$ | Dissociation constant of Calcium to CaM | $5\ \mu M$ | (10) |
| $K_{K\cdot pP}$ | Dissociation constant of PP to CaMKII | $1000\ \mu M$ | Chosen $\gg K_{pK\cdot P}$ for effective dephosphorylation |
| $K_{K\cdot ATP}$ | Dissociation constant of ATP to CaMKII | $10\ \mu M$ | (34) |
| $K_{K\cdot C_4}$ | Dissociation constant of CaM to CaMKII | $0.1\ \mu M$ | $0.045 - 5.9\ \mu M$ (14, 35) |
| $K_{K\cdot C_4\cdot ATP}$ | Dissociation constant of ATP to CaMKII | $10\ \mu M$ | Chosen equal to $K_{K\cdot ATP}$ as binding is independent |
| $K_{K\cdot ATP\cdot C_4}$ | Dissociation constant of CaM to CaMKII | $0.1\ \mu M$ | Chosen equal to $K_{K\cdot C_4}$ as binding is independent |
| $K_{pK\cdot P}$ | Dissociation constant of PP to pCaMKII | $0.001\ \mu M$ | $K_{pK\cdot P}/K_{pK\cdot C_4} \sim 1$ causing a Michaelis constant $\sim 1\mu M$ (19, 36) |
| $K_{pK\cdot ADP}$ | Dissociation constant of ADP to pCaMKII | $1\ \mu M$ | Chosen $\ll K_{K\cdot ATP}$ for phosphorylation |
| $K_{pK\cdot C_4}$ | Dissociation constant of CaM to pCaMKII | $0.001\ \mu M$ | Chosen as $K_{K\cdot C_4}/100$ due to "CaM trapping" (14) |
| $K_{pK\cdot C_4\cdot ADP}$ | Dissociation constant of ADP to pCaMKII | $1\ \mu M$ | Chosen equal to $K_{pK\cdot ADP}$ as binding is independent |
| $K_{pK\cdot ADP\cdot C_4}$ | Dissociation constant of CaM to pCaMKII | $0.001\ \mu M$ | Chosen equal to $K_{pK\cdot C_4}$ as binding is independent |
| $\alpha_K$ | Reaction rate of self-phosphorylation | $5\cdot 10^{-6}\ s^{-1}$ | Chosen $\ll \alpha_{K,NP}$ |
| $\beta_K$ | Reversal reaction rate of self-phosphorylation | $5\cdot 10^{-11}\ s^{-1}$ | Chosen $\ll \alpha_K$ |
| $\alpha_{KC}$ | Reaction rate of self-phosphorylation | $5\cdot 10^{-6}\ s^{-1}$ | Chosen $\ll \alpha_{K,NC}$ |
| $\beta_{KC}$ | Reversal reaction rate of self-phosphorylation | $5\cdot 10^{-11}\ s^{-1}$ | Chosen $\ll \alpha_{KC}$ |
| $\alpha_P$ | Reaction rate of dephosphorylation | $2.5\ s^{-1}$ | Modified from $2\ s^{-1}$ in (19) |
| $\beta_P$ | Reversal reaction rate of dephosphorylation | $5\cdot 10^{-6}\ s^{-1}$ | Chosen $\ll \alpha_P$ |
| $\alpha_{K,NC}$ | Reaction rate of neighboring phosphorylation | $10\ s^{-1}$ | Modified from $12.6\ s^{-1}$ in (37) |
| $\beta_{K,NC}$ | Reversal reaction rate of neighboring phosphorylation | $5\cdot 10^{-6}\ s^{-1}$ | Chosen $\ll \alpha_{K,NC}$ |
| $\alpha_{K,NP}$ | Reaction rate of neighboring phosphorylation | $10\ s^{-1}$ | Modified from $12.6\ s^{-1}$ in (37) |
| $\beta_{K,NP}$ | Reversal reaction rate of neighboring phosphorylation | $5\cdot 10^{-6}\ s^{-1}$ | Chosen $\ll \alpha_{K,NP}$ |
| $\mu$ | Reaction rate of opening | $1.25\ s^{-1}$ | $\nu/\mu \in [10^{-2}, 10^2]$ (20) |
| $\nu$ | Reaction rate of closing | $1\ s^{-1}$ | Modified from $0.63\ s^{-1}$ in (19) |
| $\alpha_P^{(\text{auto})}$ | Reaction rate of auto-dephosphorylation | $0.25\ s^{-1}$ | See text |
| $K_{\text{m50, pP}}$ | Michalis-Menton coefficient of auto-dephosphorylation | $1\ \mu M$ | See text |
| $\alpha_{PC}$ | Reaction rate of calcium-dependent dephosphorylation | $0.4\ s^{-1}$ | See text |
| $r_{\text{base}}$ | Baseline dephosphorylation rate | $5\cdot 10^{-5}\ s^{-1}$ | See text |
| $[P]_b$ | Baseline activity of phosphatase | $1\ \mu M$ | See text |
| $A_{tot}$ | Number of total available AMPAR | 80 | (25) |
| $c_1$ | Scaling factor | 1 | (9) |
| $c_2$ | Scaling factor | 1 | (9) |
| $c_3$ | Rate constant independent from CaMKII activity | $0.05\ s^{-1}$ | Fitting EPSP modification in (1, 28) |
| $c_4$ | Rate constant independent from PP activity | $0.15\ s^{-1}$ | Fitting EPSP modification in (1, 28) |

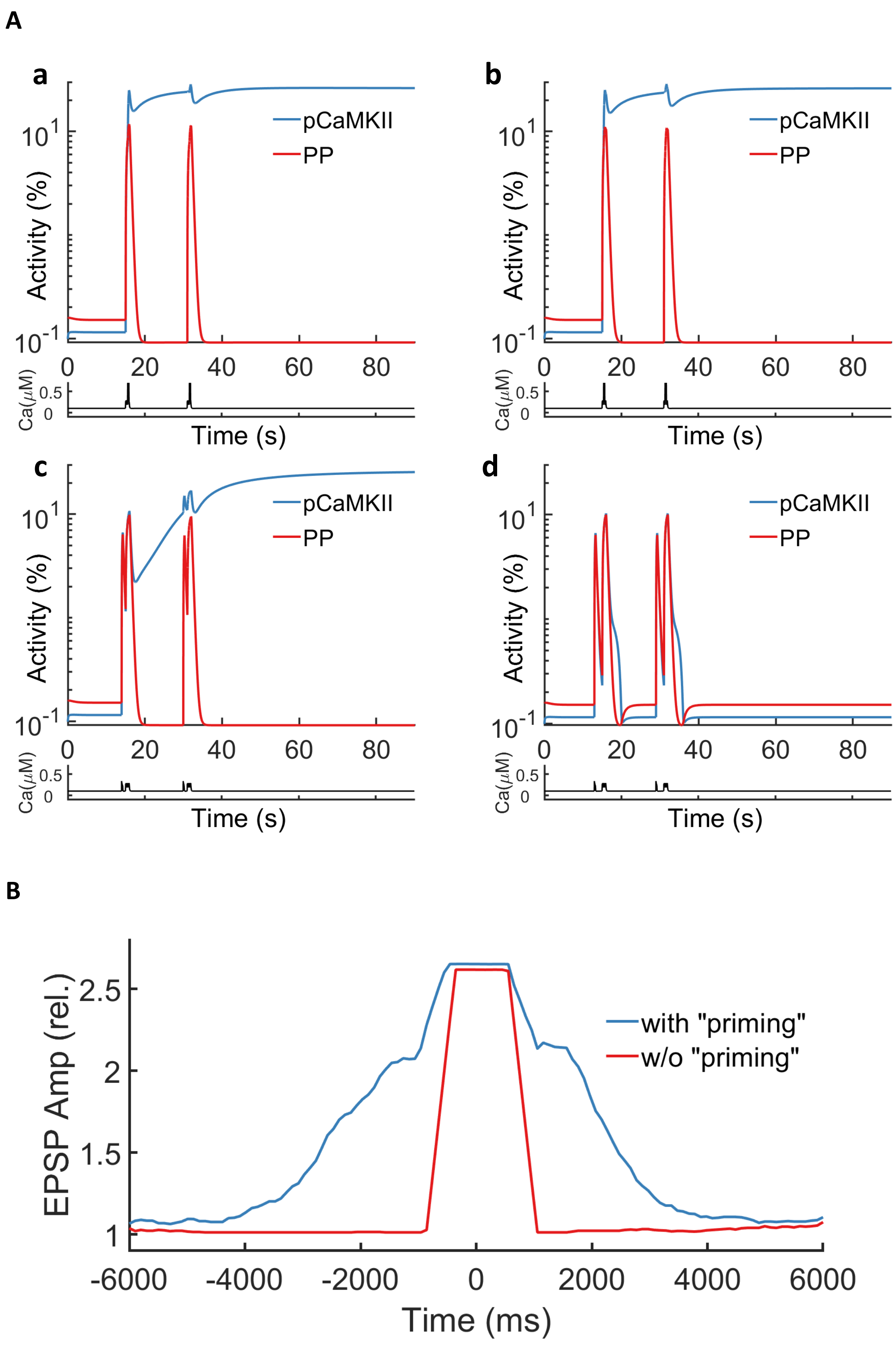


**Fig. S1.** More Results on Synaptic Modification in Response to BTSP protocols. (A) Temporal evolution of Calcium, PP1 activity, and phosphorylated CaMKII activity level during BTSP stimulation protocols. Four different scenarios are shown, with $\Delta t = -100$ ms (a), $\Delta t = 100$ ms (b), $\Delta t = 1500$ ms (c), and $\Delta t = 2500$ ms (d). Two pairs of pre-synaptic spike trains and a plateau potential (15s interval) evoke LTP with all the $\Delta t$ shown here, except $\Delta t = 2500$ ms. (B) Synaptic modifications in response to BTSP protocols, with and without DC depolarization. With depolarization of 42pA, BTSP protocols with $\Delta t$ between $-4500$ and $4000$ ms evoke LTP, and $\Delta t$ beyond that leads to almost no change (same data as in Fig.5C). However, only $\Delta t$ between $-800$ and $1000$ ms can induce LTP when DC depolarization is not presented.

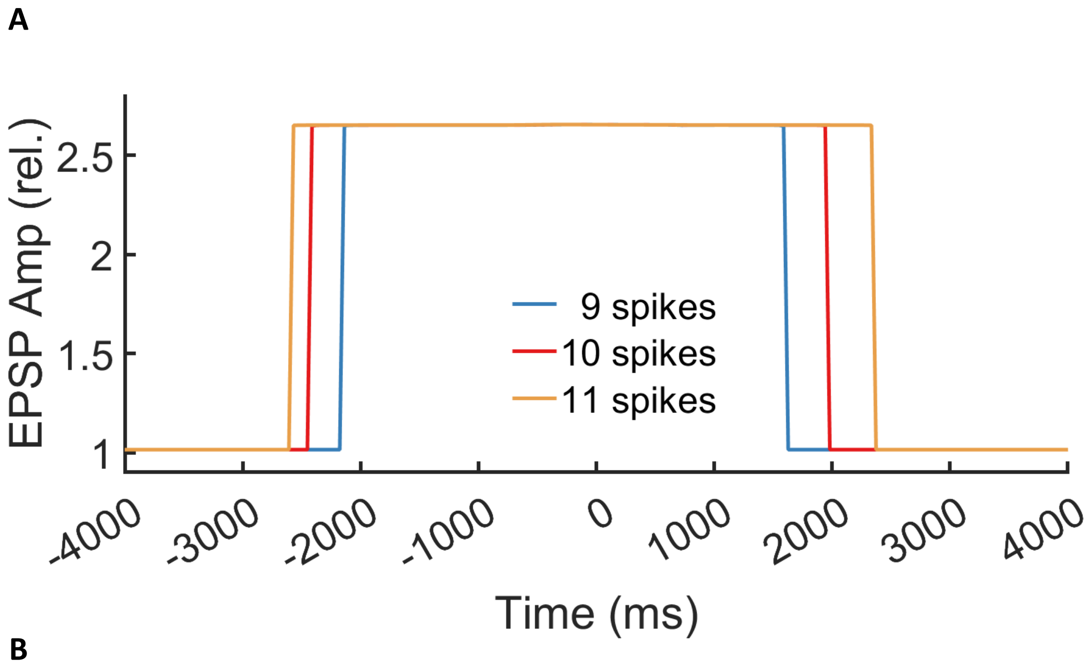


**Fig. S2.** Synaptic Modification in Response to variations of BTSP protocols. (A) Synaptic modifications in response to BTSP protocols, with varying numbers of spikes in the pre-synaptic spike trains. The range of $\Delta t$ capable of inducing LTP is $-2100$ to $1600$ ms when there are 9 spikes in one pre-synaptic spike train (blue curve), $-2400$ to $1950$ ms for 10 spikes (red curve, same data as in Fig.5C) and $-2550$ to $2350$ ms for 11 spikes (orange curve). (B) Synaptic modifications in response to BTSP protocols, with varying amplitude of plateau potential. The range of $\Delta t$ capable of inducing LTP is $-1800$ to $1600$ ms when the plateau potential is initiated by a 3500pA current injection (blue curve), $-2400$ to $1950$ ms for 4000pA (red curve, same data as in Fig.5C) and $-2950$ to $2250$ ms for 4500pA (orange curve).

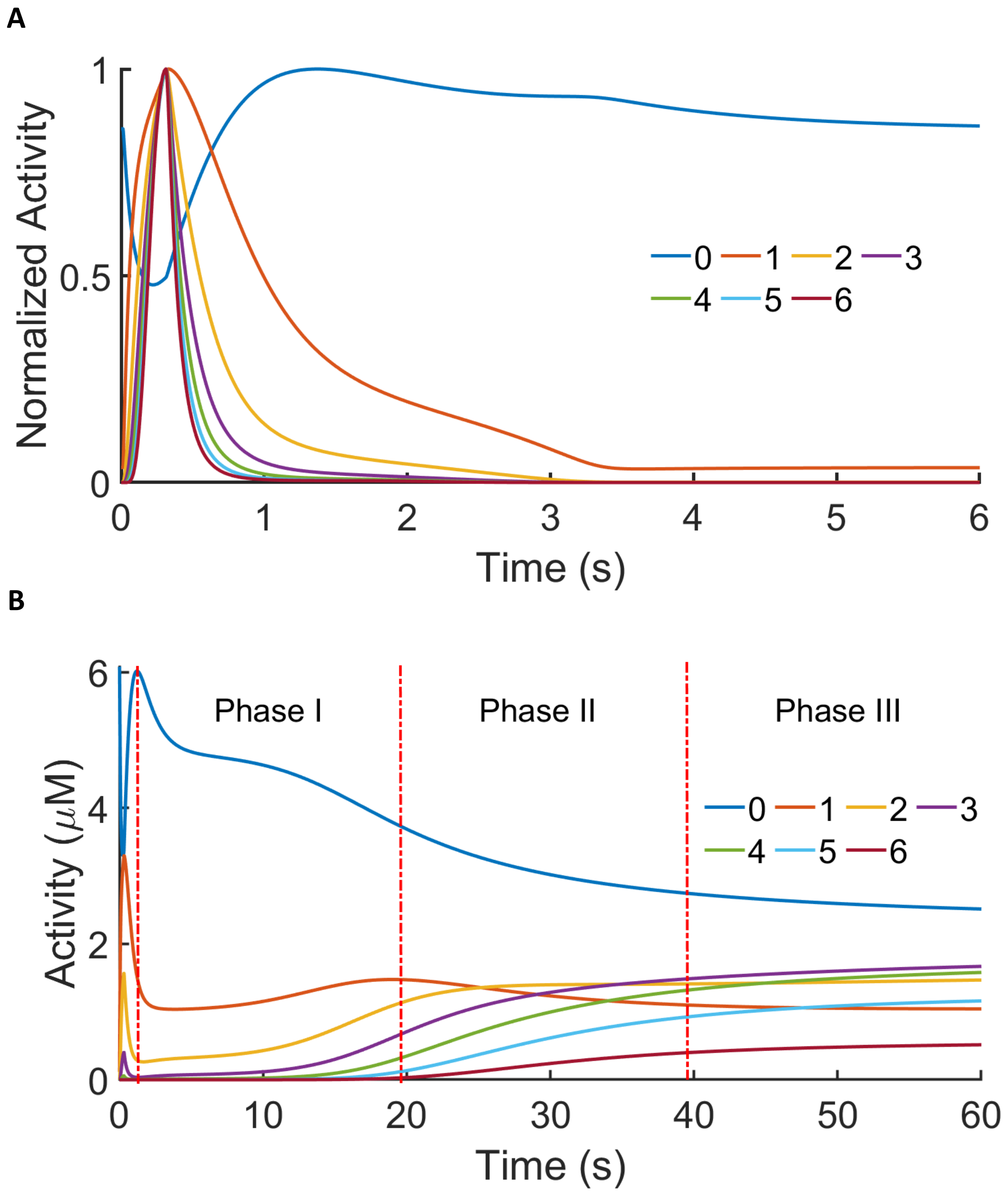


**Fig. S3.** Temporal Evolution of CaMKII Activity of Different Phosphorylation Levels. (A) Temporal evolution of CaMKII activity (normalized) of different phosphorylation levels in response to a plateau potential. The plateau potential is triggered by a 5000pA current injection, lasting for 300ms, starting at $t = 0$ second, and a sustained 42pA depolarization throughout. The trace labeled 'k' indicates the activity of CaMKII with k subunits phosphorylated. (B) Temporal evolution of CaMKII of different phosphorylation levels in response to a plateau potential with a perturbation on open CaMKII activity. The same stimulus is used as in A, with an additional increase in the concentration of CaMKII with no phosphorylated subunits by $0.8\mu$M (and a corresponding decrease in closed CaMKII by $0.8\mu$ M) at the beginning of the plateau potential. The CaMKII activity traces can be divided into three phases: (I) plateau, (II) increasing, and (III) stabilization.

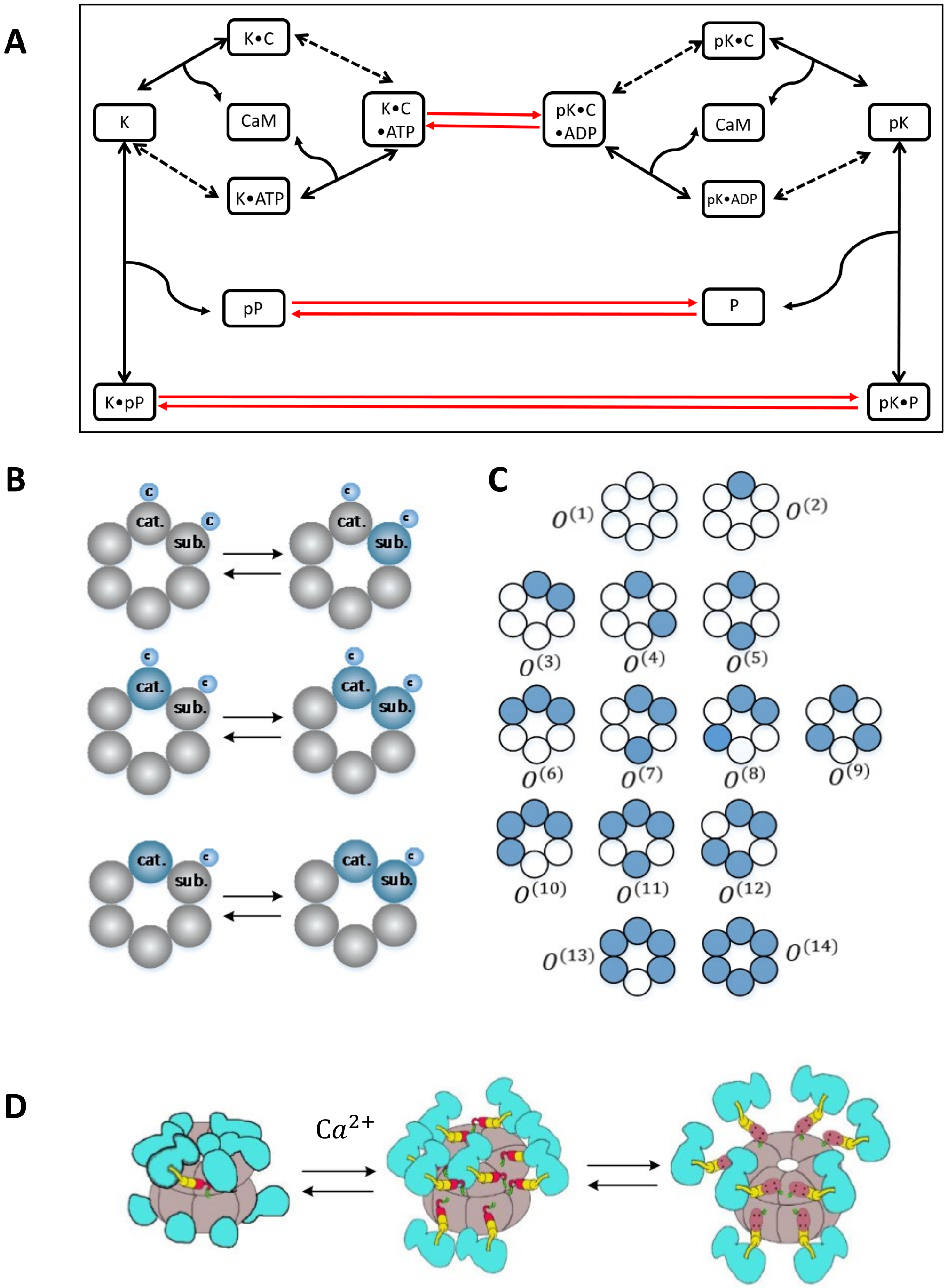


**Fig. S4.** Modeling CaMKII Phosphorylation and Conformational Changes. (A) The reactions for a single CaMKII subunit. The black arrows represent the binding and disassociation of CaMKII subunits with PP, ATP/ADP, and CaM, modeled as quasi-equilibrium. The red arrows indicate the phosphorylation and dephosphorylation of CaMKII and PP, modeled as ordinary differential equations. The dashed black arrows represent the binding of phosphorylated/unphosphorylated CaMKII subunits with ADP/ATP, respectively. (B) The three pathways for the neighboring phosphorylation. The catalytic and substrate subunit are marked as "cat." and "sub.", respectively. Unlabeled subunits do not affect the phosphorylation of the substrate subunit. The substrate subunit must be bound with ATP and CaM for the phosphorylation to occur. The catalytic subunit can be unphosphorylated but bound with CaM (a) or phosphorylated, either with (b) or without (c) CaM bounded. (Adapted from (Fig. 1 in (2)) (C) Different states of a single CaMKII ring of 6 subunits. Each subunit can be either phosphorylated (blue) or unphosphorylated (gray). A total of 14 conformations of one subunit can be found under rotation invariance, marked as $O^{(i)}, 1 \leq i \leq 14$. (D) The docking and undocking of CaMKII. CaMKII exists in both compact ($C$) and extended auto-inhibited states ($O^{(i)}$) and can freely transit between the two states with a calcium-dependent rate. CaMKII in the compact state $C$ must become extended $O^{(1)}$ before it can be phosphorylated. (Adapted from Fig.5 in (21).)

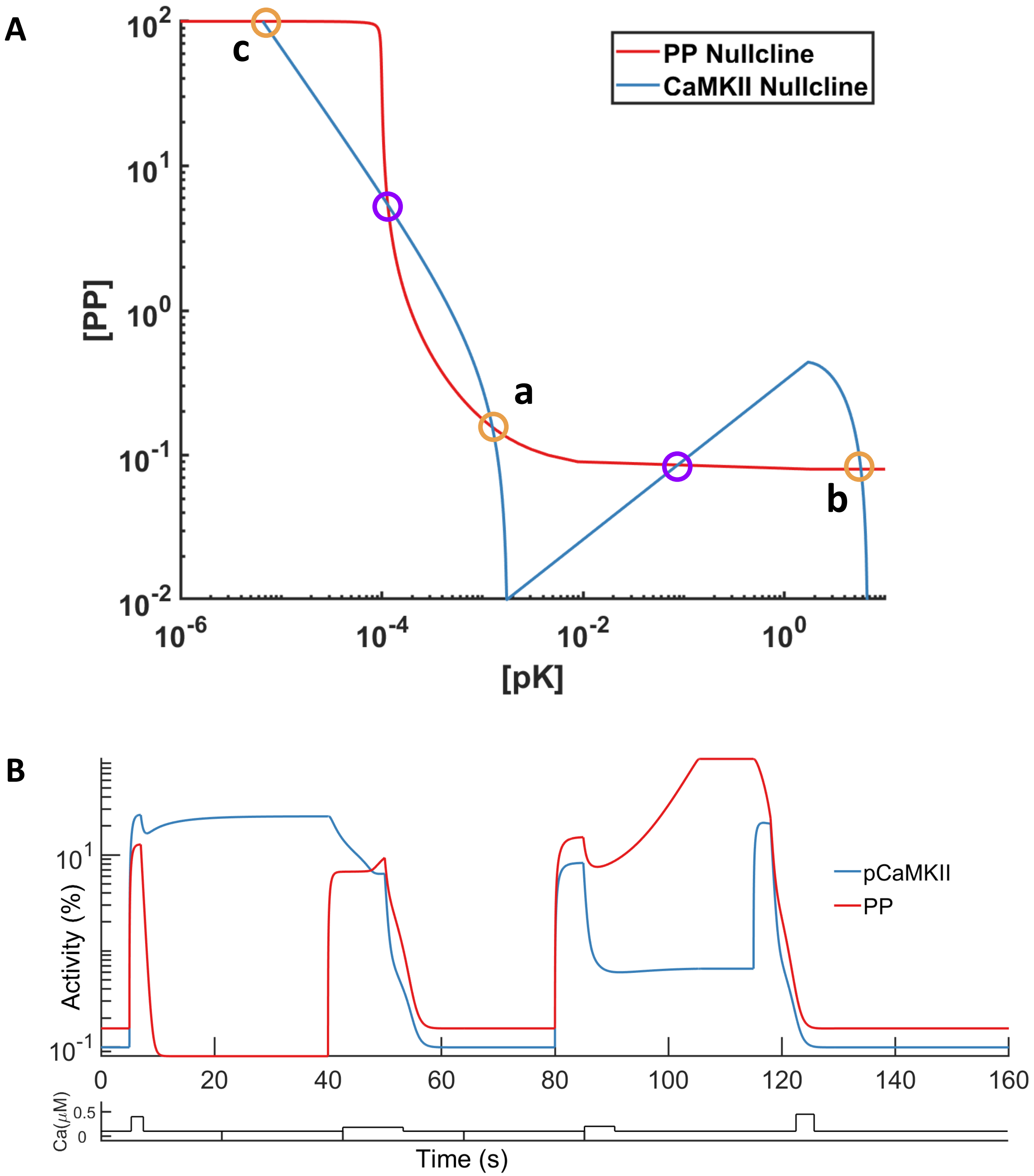


**Fig. S5.** Dynamics of CaMKII and PP interaction. (A) The phase plane of the model of CaMKII and PP Interaction. Blue and red curves indicate nullclines for phosphorylated CaMKII and active PP, respectively. The purple circles mark the two unstable steady states, while the orange circles indicate the three stable steady states, corresponding to basal (a), LTP(b), and LTD (c) states. (B) Phosphorylated CaMKII and active PP levels during the reversals of LTP and LTD. A calcium stimulus protocol similar to (9) is applied with four pulses. The first calcium pulse stimulus ($0.3\mu M$ for 2 s) induces the system from the basal to the LTP state. The potentiated state is reversed (depotentiation) by the second pulse ($0.08\mu M$ for 10 s). LTD is induced by the third pulse ($0.1\mu M$ for 5 s) and then reversed (dedepression) by the fourth stimulus ($0.35\mu M$ for 3 s).


Guanchun Li, David W. McLaughlin, Charles S. Peskin

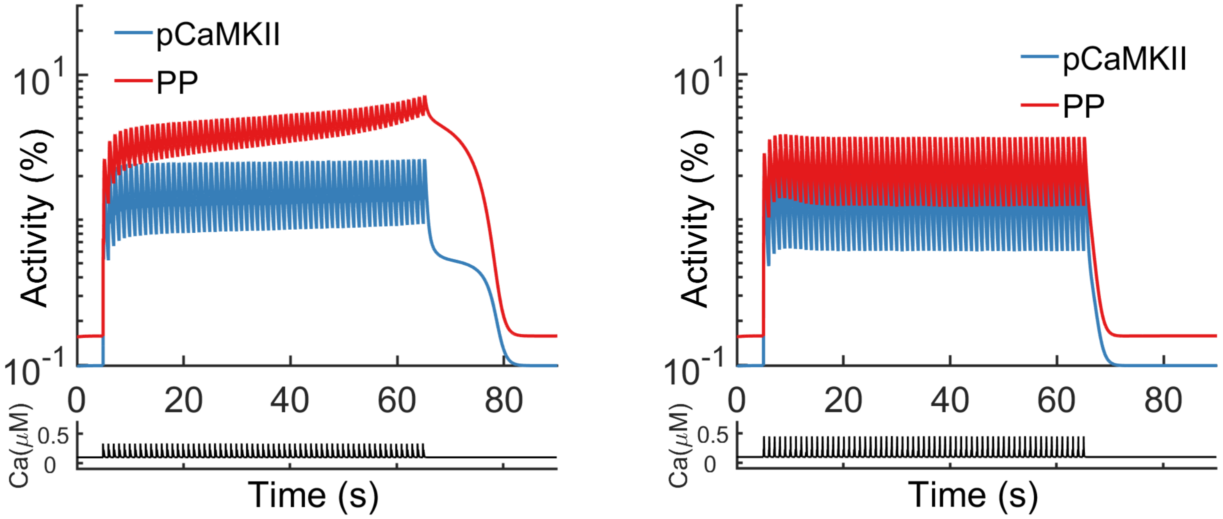


**Fig. S6.** More results on Synaptic Modification in Response to STDP protocols. Temporal evolution of Calcium, PP1 activity, and phosphorylated CaMKII activity level during STDP stimulation protocols with $dt = 100$ ms (left panel) and $dt = -100$ ms (right panel). 60 pairs of pre-synaptic and post-synaptic spikes in 1Hz leads to no change when $dt = \pm 100$ ms.

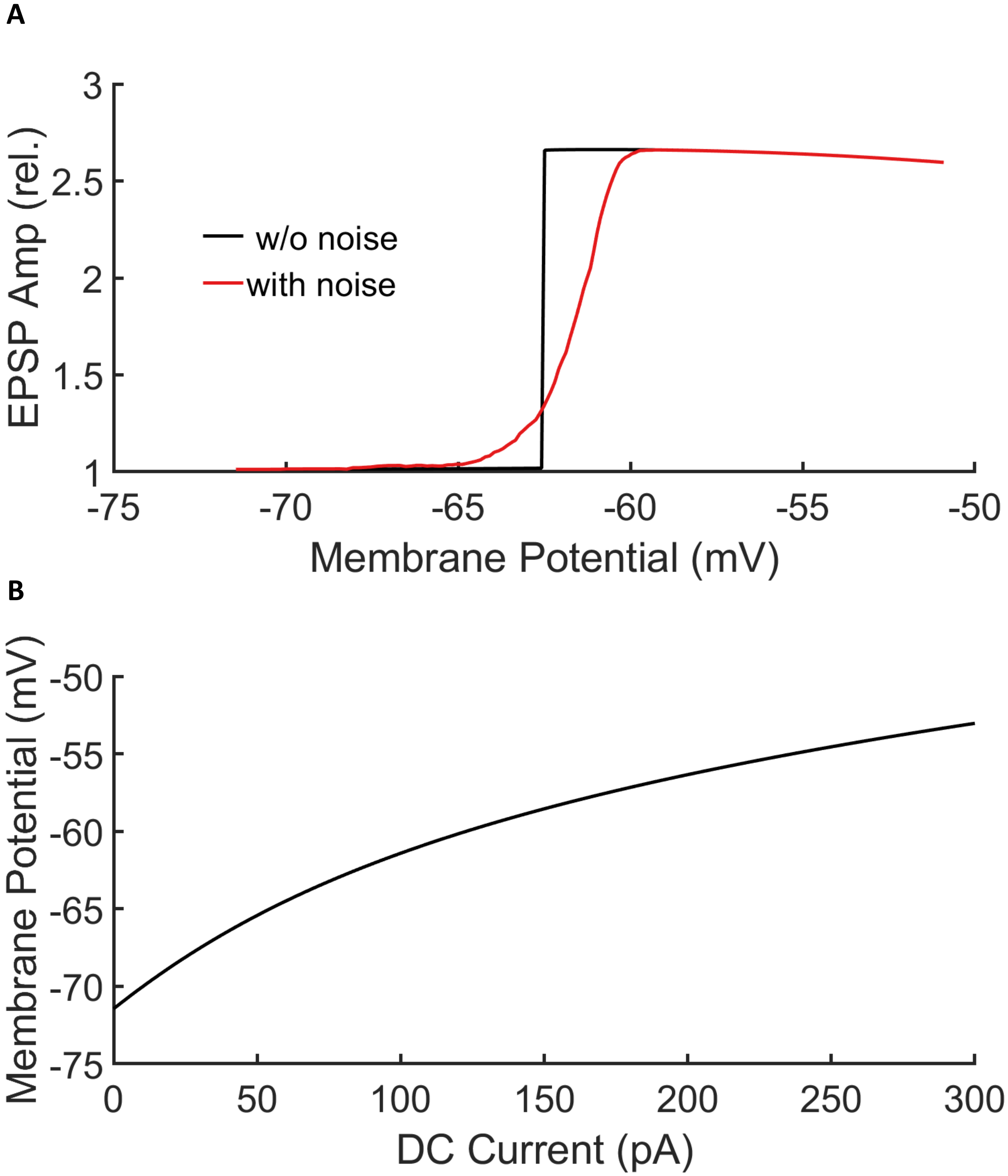


**Fig. S7.** More results on Synaptic Modification in Response to Voltage-Dependent Learning. (A) Synaptic modifications in response to voltage-dependent "one-shot" learning with varying baseline membrane potential. As the baseline membrane potential is adjusted through DC depolarization, a clear transition to learning is seen at -62.5mV in the deterministic setting (black curve), and a similar transition is observed within the range of $-65 \sim -60$ mV for the stochastic setting (with the half height of the transition at $-61.5$ mV). (B) The relationship between baseline membrane potential and DC current amplitude. The baseline membrane potential is shown to be a monotonic function of subthreshold DC current amplitude.

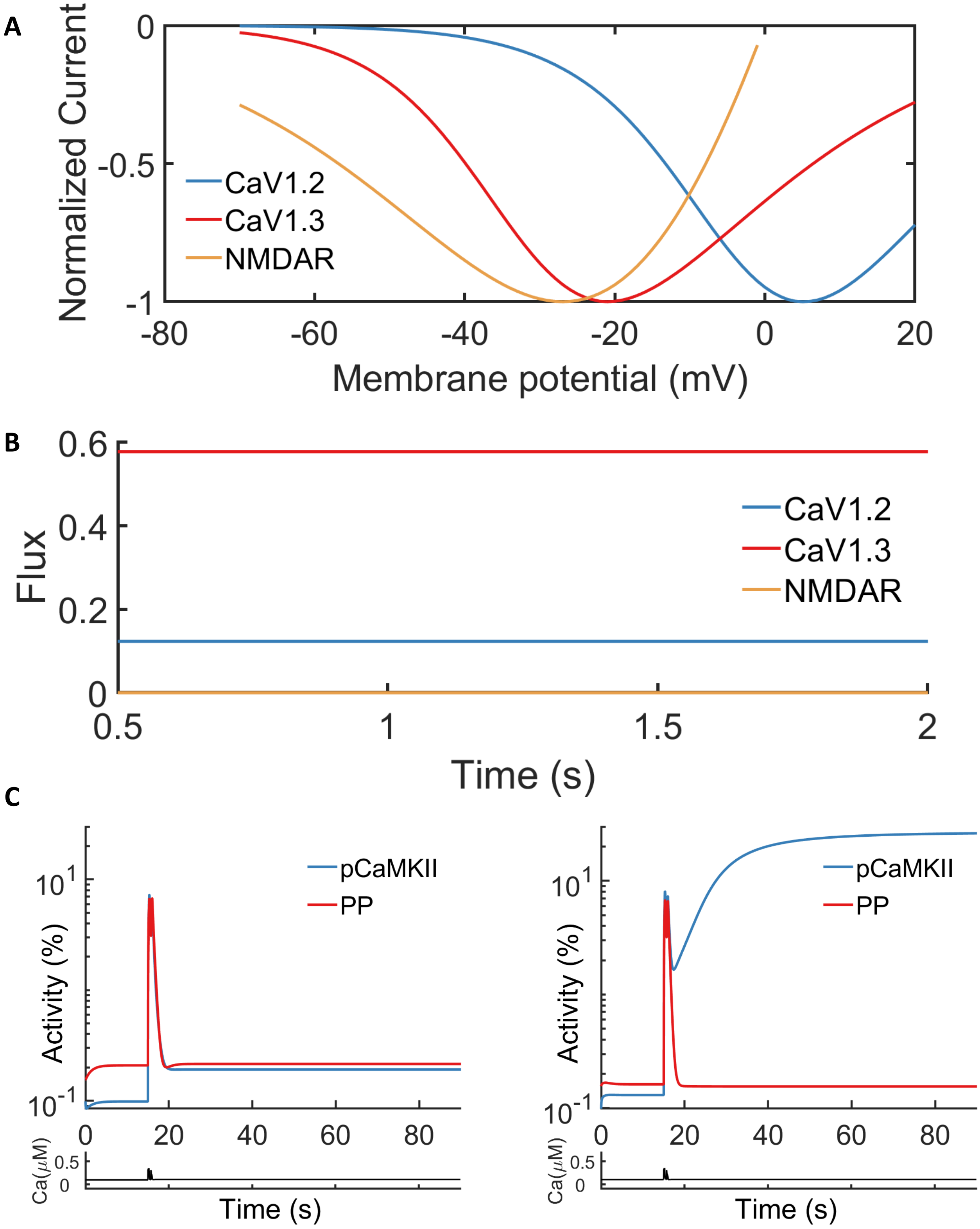


**Fig. S8.** Voltage Dependency of Calcium Channels in "one-shot" learning. (A) Voltage-dependency of NMDAR, CaV1.2, and CaV1.3 channels in the model. The curves show the peak current (normalized) flowing through the corresponding channels when the membrane potential is increased to the given value as a step function starting from the resting membrane potential. The results are similar to the experimental measurements (see Fig. 5b, (38)). (B) Temporal evolution of Calcium flux through NMDAR, CaV1.2, and CaV1.3 channels with subthreshold depolarization. The traces show the calcium flux through each channel type during a DC depolarization of 42pA. CaV1.3 channels contribute the majority (more than 80%) of calcium flux during the DC depolarization. (C) Temporal evolution of Calcium, PP1, and phosphorylated CaMKII activity level during the "one-shot" learning protocol, with CaV1.2 or CaV1.3 channels blocked. The stimulus protocol is the same as Fig.3B except that a higher DC depolarization (120pA) is applied. Blocking CaV1.3 channels (left panel) abolishes LTP while blocking CaV1.2 channels (right panel) does not affect LTP.

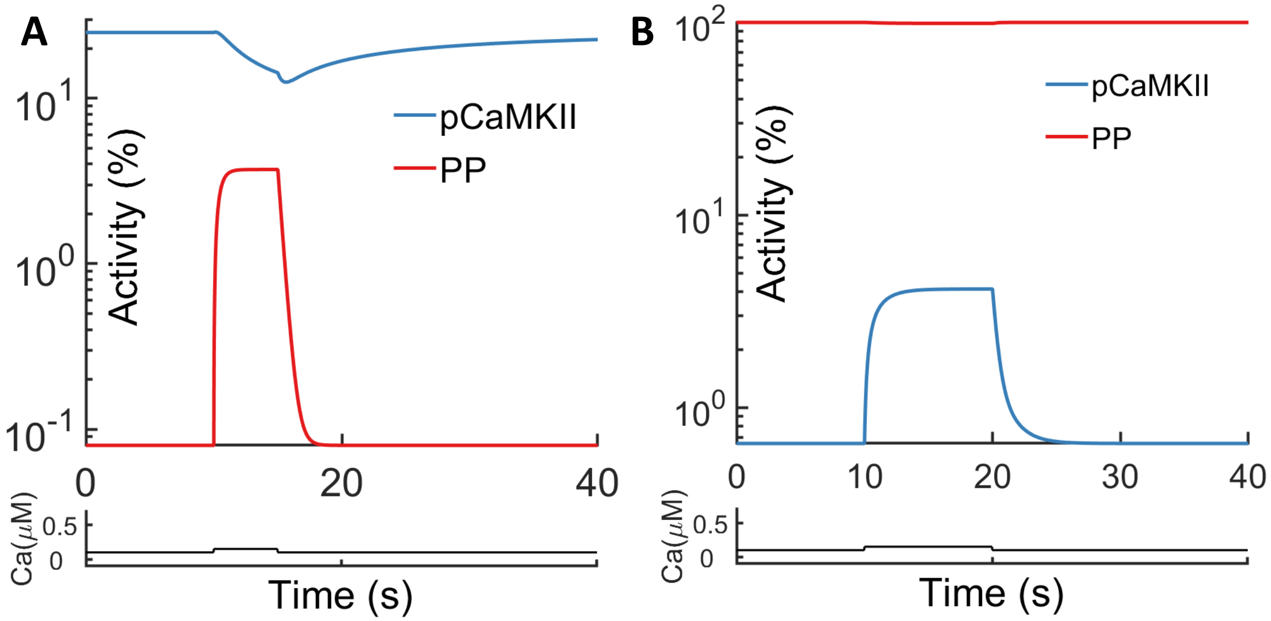


**Fig. S9.** Dynamics of CaMKII and PP Interaction. (A) Phosphorylated CaMKII and active PP levels during perturbation in the LTP state. Initially at the LTP steady state, the system remains in the LTP state after the perturbation, a calcium pulse stimulus ($0.05\mu M$ for 10 seconds) injected at $t = 10s$. (B) Phosphorylated CaMKII and active PP levels during perturbation in the LTD state. Initially at the LTD steady state, the system remains in the LTD state after the perturbation, a calcium pulse stimulus ($0.05\mu M$ for 10 seconds) injected at $t = 10s$.
